# Recording of cellular physiological histories along optically readable self-assembling protein chains

**DOI:** 10.1101/2021.10.13.464006

**Authors:** Changyang Linghu, Bobae An, Monika Shpokayte, Orhan T. Celiker, Nava Shmoel, Chi Zhang, Won Min Park, Steve Ramirez, Edward S. Boyden

## Abstract

Observing cellular physiological histories is key to understanding normal and disease-related processes, but longitudinal imaging is laborious and equipment-intensive. A tantalizing possibility is that cells could record such histories in the form of digital biological information within themselves, for later high-throughput readout. Here we show that this concept can be realized through information storage in the form of growing protein chains made out of multiple self-assembling subunits bearing different labels, each corresponding to a different cellular state or function, so that the physiological history of the cell can be visually read out along the chain of proteins. Conveniently, such protein chains are fully genetically encoded, and easily readable with simple, conventional optical microscopy techniques, compatible with visualization of cellular shape and molecular content. We use such expression recording islands (XRIs) to record gene expression timecourse downstream of pharmacological and physiological stimuli, in cultured neurons and in living mouse brain.

## Introduction

Reading out biological signals and processes that take place over time, in living cells, intact organs, and organisms, is essential to advancing biological research, both basic science and translationally oriented. The imaging of genetically encoded fluorescent signal reporters, for example, enables specific biological activities to be monitored in real time in living cells^1^. However, long-term live imaging is laborious and equipment intensive, because a single microscope often has to be monopolized for the duration of the experiment, and furthermore the number of cells that can be observed is limited by the performance of live imaging methods, which are not as scalable as fixed-tissue imaging methods, which can benefit from sectioning, clearing, expansion, and other techniques that improve the number of cells that can be surveyed, the resolution, and the number of signals that can be analyzed at once^2–4^. Snapshot methods, that perform RNA FISH^5^ or protein immunostaining^6^, can enable one (and sometimes two) timepoints of a physiological signal to be inferred in fixed cells, but cannot support continuous recording of physiological signals for later fixed-cell readout. Nevertheless, these methods allow biological information readout over very large spatial scales, even entire mammalian brains, because the fixed cells or tissues can be scalably imaged thanks to the aforementioned post-preservation tissue processing strategies.

In principle, if biological information could be recorded by cells and stored, digitally, within their own cellular volumes, for later readout after cell fixation or other preservation strategies, it could be possible to get the best of both worlds – recording of continuous time histories of physiological signals, followed by scalable fixed tissue signal history readout. Several studies have proposed, and demonstrated, the recording of cellular histories into nucleic acid form, for readout through high-throughput nucleic acid sequencing that requires the cells or tissues to be dissociated and/or lysed^7–16^. But often one wants to consider such physiological histories in intact cellular, tissue, or organ contexts – hence the popularity of imaging as a strategy throughout biology. We discovered that it is possible to record biological information along growing protein chains made out of fully genetically encoded self-assembling proteins which bear different labels that encode for different cellular states or functions. While the cell is alive, the self-assembling proteins (bearing labels) are added constantly to the growing chain, enabling continuous recording of the presence of the different label-bearing proteins that are available (**Figure 1a,b**). For example, if at a certain point in time, proteins with one label are common, and proteins with another label are rare, the part of the chain that is growing at the current moment in time, will have more of the former label than the latter. In other words, the local density of labels will favor the first label over the second, even if the labels are independent and being added at a constant rate. When the experiment is done, the chain of proteins can be read out by ordinary immunostaining and imaging, after cell or tissue fixation.

**Figure 1.**
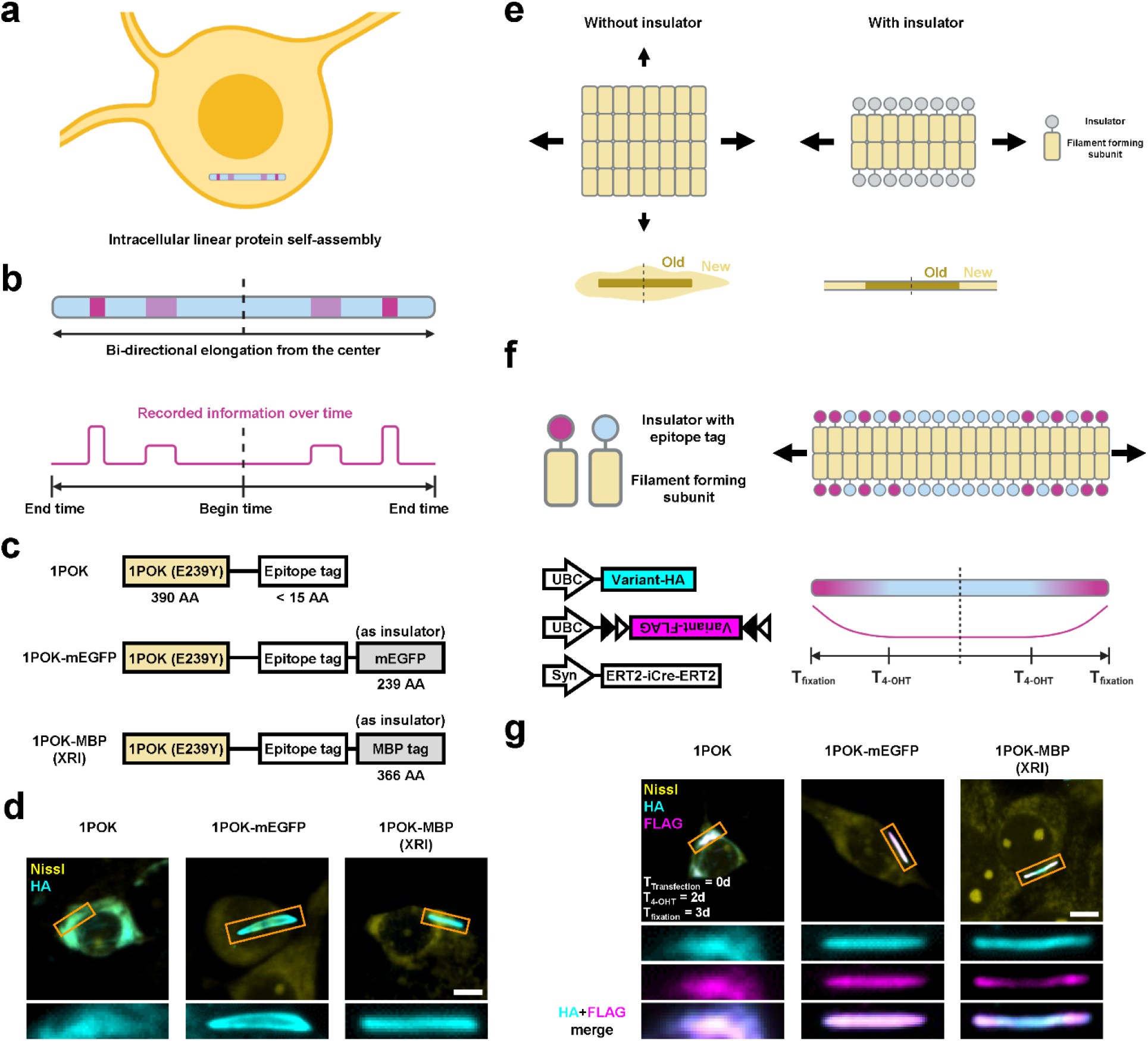
Concept and development of linear protein-based cellular physiology recording devices. (**a**) Schematic of intracellular linear protein self-assembly. (**b**) Schematic of bi-directional elongating intracellular linear protein self-assembly for encoding, storing, and reading out biological information. Blue shading, components on the self-assembly whose expression is constitutive over time; red shading, components on the self-assembly whose expression is dependent on biological events of interest over time; red line, density along the self-assembly of the components whose expression is dependent on biological events of interest over time. (**c**) Schematic of variants of self-assembling proteins. 1POK (E239Y), a filament-forming self-assembling protein; mEGFP, monomeric enhanced green fluorescent protein; MBP tag, maltose binding protein tag; AA, amino acid; XRI, the variant that was selected as the XRI design throughout this paper (see **Supplementary Table 1** for sequences of the motifs; see **Supplementary Table 2** for all tested constructs). (**d**) Representative confocal images of cultured mouse hippocampal neurons expressing self-assembling protein variants with the epitope tag HA, taken after fixation, Nissl staining, and immunostaining against the HA tag. Scale bar, 5 μm throughout this figure. Rectangular panels at the bottom, enlarged views of regions marked in orange rectangles in the top row of square panels. (**e**) Schematic of protein self-assemblies without (left) and with (right) a insulator component fused to each of the filament forming subunits. Arrows with different sizes, growth directions of protein self-assemblies, with arrow sizes indicating growth rates; old, subunits that bound to the protein self-assembly earlier; new, subunits that are binding to the protein self-assembly currently. (**f**) Schematic of the protein self-assembly and the constructs in the chemically induced gene expression experiment. Variant-HA, self-assembling protein variant (1POK, 1POK-mEGFP, or 1POK-MBP) with the epitope tag HA; Variant-FLAG, self-assembling protein variant with the epitope tag FLAG; UBC, human ubiquitin promoter; Syn, human synapsin promoter; black and white triangles, lox sites in the FLEX construct; 4-OHT, 4-hydroxytamoxifen; T_4-OHT_, the time when cells are treated with 4-OHT; T_fixation_, the time when cells are fixed. (**g**) Representative confocal images of cultured mouse hippocampal neurons expressing constructs (shown in bottom left of **f**), taken after fixation, Nissl staining, and immunostaining against the HA tag and the FLAG tag. T_transfection_, the time when the constructs are delivered to cells via DNA transfection. Three rows of rectangular panels at the bottom, enlarged views of regions marked in orange rectangles in the top row of square panels.

We show that this expression recording island (XRI) strategy can be used for long-term recording of gene expression time course, with single-cell precision, across cell populations. Because the linear protein assembly grows continuously over time, it acts like a molecular tape recorder that preserves the temporal order of the protein monomers made available by the cell depending on the cell’s current state or function. For example, if protein monomers with the epitope tag ‘A’ are steadily expressed by the cell, and the expression of protein monomers with the epitope tag ‘B’ is increased by, say, a neural activity dependent promoter, then the neural activity dependent event will result in permanent storage of the activity record in the order of the epitope tags along the growing protein chain, enabling later readout via immunostaining against tags ‘A’ and ‘B’, followed by standard imaging. We applied XRIs to perform 4-day recordings of, amongst other things, c-fos promoter-driven gene expression in cultured mouse hippocampal neurons after depolarization, and also showed that pharmacological modulation of gene expression histories in the living mouse brain could also be read out post hoc.

## Results

We first set out to test if human-designed proteins known to self-assemble into filaments, could be coaxed to reliably form continuously growing linear chains in cultured mammalian cells. We fused 14 human-designed filament-forming proteins (previously characterized in buffers, bacteria, and yeast) to a short epitope tag (HA, for immunofluorescence imaging after protein expression and cell fixation) and expressed them in primary cultures of mouse hippocampal neurons (see **Supplementary Table 1** for sequences of the motifs; see **Supplementary Table 2** for all tested constructs). Upon immunofluorescence staining, followed by imaging under confocal microscopy, two filament-forming proteins produced clear and stable fiber-like structures in the cytosol: 1POK (from Levy et al.)^17^ (**Figure 1c,d**) and DHF40 (from Baker et al.)^18^ (**Supplementary Figure 1a**). The rest of the proteins produced unstructured aggregates, high non-assembly background, and/or punctum-like structures in neurons (see **Supplementary Figure 1b** for example; see **Supplementary Table 2** for complete screening results). However, both filament-forming proteins also produced unstructured aggregates of protein in the cytosol. DHF40 showed a higher immunofluorescence background in cytosolic areas, which did not correspond either to fiber-like structures or unstructured aggregates, than did 1POK, suggesting DHF40 had a higher level of free-floating protein monomer that did not bind to the protein assembly, than did 1POK. Due to the lower immunofluorescence background, we selected 1POK as the filament forming protein for further engineering in this study.

Because linear protein assembly would enable useful information encoding that could then be easily read out, we next performed protein engineering on 1POK to reduce the unstructured aggregates in cells. We reasoned that unstructured aggregates could be present due to unwanted lateral growth (**Figure 1e, left**), as opposed to the longitudinal growth that would result in linear information encoding, and that reducing such lateral growth would discourage the formation of unstructured aggregates and thus encourage fiber-like linear protein assembly (**Figure 1e, right**). We hypothesized that, by fusing a filament “insulator” component to the lateral edge of the filament forming monomer, unwanted lateral binding and growth of the protein assembly would be sterically blocked. We fused highly monomeric proteins that are widely used in bioengineering, mEGFP^19^ (a green fluorescent protein) and maltose binding protein (MBP tag; an *E. coli* protein commonly used as a solubility tag for recombinant protein expression in mammalian^20^ and non-mammalian^21^ cells) to 1POK as insulators, together with the short epitope tag HA (**Figure 1c**). We chose monomeric proteins as insulators because we reasoned that any homo-oligomeric binding of non-monomeric proteins might encourage, rather than halt, unwanted lateral binding and growth of the protein assembly. Expression of these variants in mouse neurons showed that both produced only fiber-like structures, without any unstructured aggregates (**Figure 1d**).

Next, we tested if the mEGFP or MBP tag-bearing variants could encode information along their linear extent while preserving temporal order of the information along their corresponding protein assemblies. If protein monomers with, say, the epitope tag HA are constantly expressing, and the expression of protein monomers with, say, the epitope tag FLAG are induced at a specific timepoint, then at that timepoint, monomers with the FLAG tag will be more common than before, and thus added at a higher rate than before, along the growing protein chain. Then, the period of time at which FLAG is expressed could be easily read out via immunostaining against both HA and FLAG tags (**Figure 1f**). We used the ERT2-iCre-ERT2 based chemically inducible Cre system^22^ to activate the expression of protein monomers with the FLAG tag, in a Cre-dependent FLEX vector, by 4-hydroxytamoxifen (4-OHT) treatment at defined times (**Figure 1f**). Co-expressing these two vectors, both driven by the constitutive human ubiquitin (UBC) promoter, with a continuously expressed HA-bearing monomer in mouse neurons via DNA transfection, and then treating the neurons with 4-OHT for 15 minutes at a time point 2 days after transfection, was followed by fixing the neurons 1 day later, followed in turn by processing for immunofluorescence. We performed this experiment for each of the three variants, 1POK, 1POK-mEGFP, and 1POK-MBP (**Figure 1g**). For the original 1POK variant without the insulator (**Figure 1g**, left), we found a high similarity between the immunofluorescence patterns of the HA tag and the FLAG tag, showing that the 1POK variant could not preserve the temporal order of the protein monomers expressed, as we had hypothesized (**Figure 1e**). For the 1POK-mEGFP variant (**Figure 1g**, middle), we also found a high similarity between the immunofluorescence patterns of the HA tag and FLAG tag. We hypothesized that this might be due to the existence of a small but non-negligible unwanted lateral growth in this variant, so that newly expressed FLAG-fused monomers coated the lateral boundaries of the entire fiber assembly, resulting in uniform immunofluorescence of the FLAG tag along the assembly. For the 1POK-MBP variant, we found the immunofluorescence of the HA tag to show a continuous intensity profile along the protein assembly (**Figure 1g**, right), while that of the FLAG tag showed higher intensity towards the two ends of the protein assembly and lower intensity towards the center of the protein assembly, a more polarized pattern. Thus, the 1POK-MBP variant showed a pattern that preserves temporal information created by the triggering of the FLAG tag at a defined point in time. We named this variant as the XRI, going forward throughout the rest of the study.

To study how accurate this XRI protein assembly could preserve time information, we again used the chemically-inducible Cre system and treated different neuron cultures expressing XRI with 4-OHT at different times after beginning of expression. To increase the efficiency of gene delivery, we used adeno-associated viruses (AAVs) to deliver the chemically-inducible Cre system and the XRI genes into cultured mouse neurons. Because the expression of AAV is slower compared to DNA transfection, we increased the expression time window from 3 days to 7 days before fixation, immunofluorescent labeling, and imaging. We divided the neuron cultures into 7 groups, and added 4-OHT treatment at 1, 2, 3, 4, 5, or 6 days after AAV transduction, or not at all (**Figure 2a-c**). We found continuous HA immunofluorescence in neurons in all groups (**Figure 2d**). We found XRI assemblies to have no FLAG immunofluorescence in neurons without 4-OHT treatment, indicating negligible leak expression of the chemically inducible Cre system (**Figure 2d**, ‘No 4-OHT’ panel). We found FLAG immunofluorescence to have strong polarized patterns (e.g., brighter at the ends than in the middle) in neurons with 4-OHT treatment on day 3, 4, 5, or 6 after AAV transduction, but not to have strongly polarized patterns in neurons with 4-OHT treatment on day 1 or 2 after AAV transduction (**Figure 2d-2e**); the HA tag showed a gentle polarization trend in the opposite direction, perhaps because the HA-bearing subunits available were landing on the growing protein chain at greater distances than before, due to the FLAG-bearing subunits having already been added. We reasoned that on 1 and 2 days after AAV transduction the XRI assemblies either did not form stably, or did form stably but with a substantial amount of lateral growth. Thus, the XRI can start reliably recording the expression time course of FLAG-bearing monomers 3 days after AAV transduction, but not 1 or 2 days after AAV transduction.

**Figure 2.**
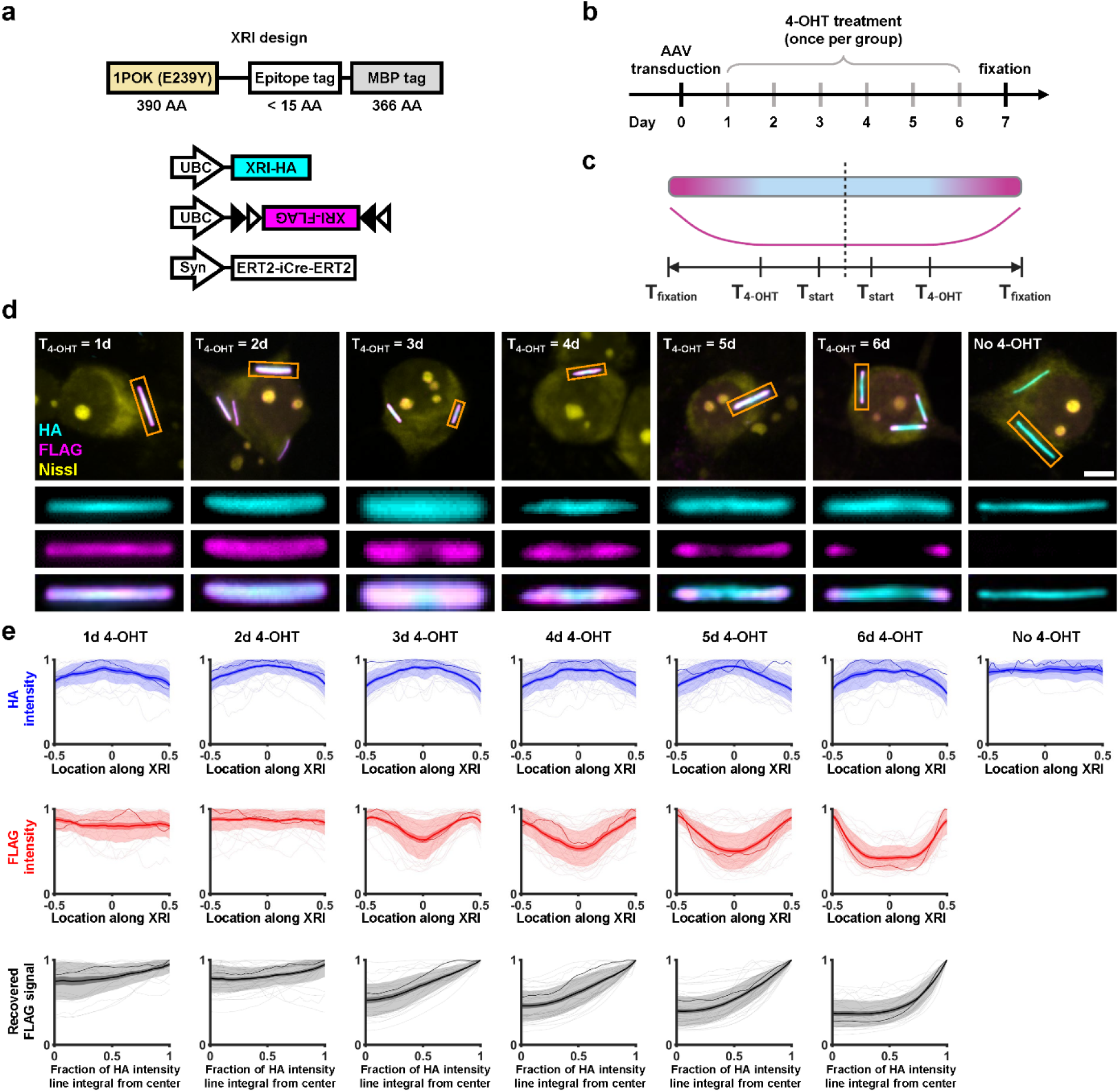
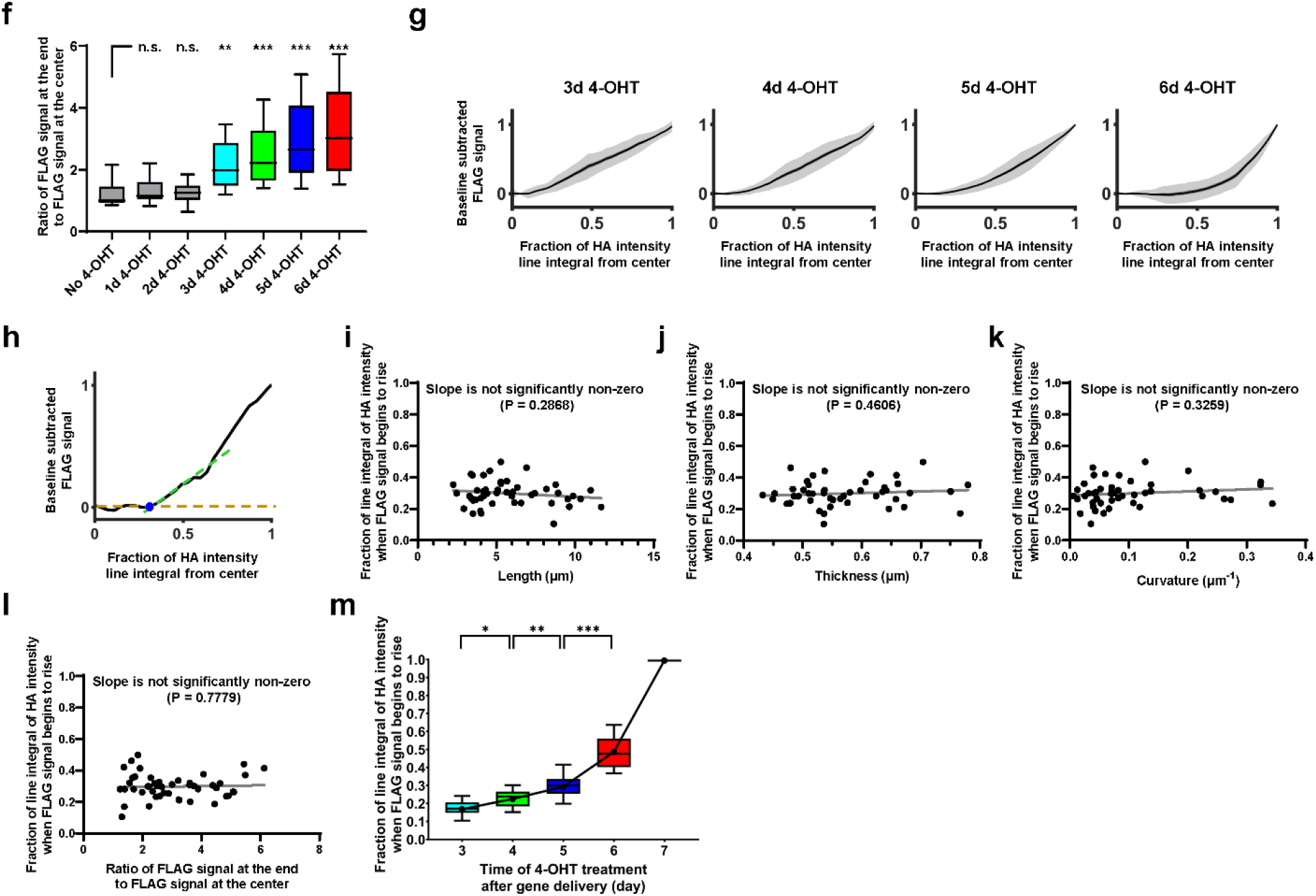
Characterization and calibration of XRIs via timed chemically induced expression. (**a-c**) Schematics of the constructs co-transduced into neurons (**a**), experiment pipeline (**b**), and expected epitope distribution along the XRI protein self-assembly (**c**) in the chemically induced gene expression experiment. XRI-HA, XRI with the epitope tag HA; XRI-FLAG, XRI with the epitope tag FLAG. The constructs were delivered to cells on day 0 via adeno-associated virus (AAV) transduction, and fixed 7 days later (T_fixation_ = 7 days). T_4-OHT_, time of 4-OHT treatment (once only per group of neurons); T_start_, the time when XRI starts recording information after gene delivery and expression of XRI (see **g**, where T_start_ is measured to be 3 days after AAV transduction). (**d**) Representative confocal images of cultured mouse hippocampal neurons expressing constructs in **a**, taken after fixation, Nissl staining, and immunostaining against HA and FLAG tags. Three rows of rectangular panels at the bottom, enlarged views of regions marked in orange rectangles in the top row of square panels. Scale bar, 5 μm. (**e**) HA intensity profile along the XRI (top row), FLAG intensity profile along the XRI (middle row), and recovered FLAG signal (by averaging the two FLAG intensity profiles from the two halves of the XRIs) plotted against the fraction of the line integral of HA intensity (a value between 0 and 1; 0 corresponds to the center of the XRI, and 1 corresponds to the end of the XRI; bottom row), from the experiment described in **a-c** (n = 21 XRIs from 13 neurons from 2 cultures for ‘1d 4-OHT’ group; n = 37 XRIs from 19 neurons from 2 cultures for ‘2d 4-OHT’ group; n = 32 XRIs from 22 neurons from 2 cultures for ‘3d 4-OHT’ group; n = 38 XRIs from 22 neurons from 2 cultures for ‘4d 4-OHT’ group; n = 47 XRIs from 32 neurons from 2 cultures for ‘5d 4-OHT’ group; n = 29 XRIs from 19 neurons from 2 cultures for ‘6d 4-OHT’ group; n = 9 XRIs from 4 neurons from 1 culture for ‘No 4-OHT’ group). Each raw trace was normalized to its peak to show relative changes before averaging. Thick centerline, mean; darker boundary in the close vicinity of the thick centerline, standard error of mean; lighter boundary, standard deviation; lighter thin lines, data from individual XRIs; darker thin line, data from the corresponding XRI in the orange rectangle in **d**. See **Supplementary Figure 2** for the detailed process flow of extracting signals from XRI assemblies. (**f**) Box plot of the ratio of the FLAG signal at the end of XRI to the FLAG signal at the center of XRI. Middle line in box plot, median; box boundary, interquartile range; whiskers, 10-90 percentile. n.s., not significant; **, P < 0.01; ***, P < 0.001; Kruskal-Wallis analysis of variance followed by post-hoc Dunn’s test with ‘No 4-OHT’ as control group. See **Supplementary Table 3** for details of statistical analysis. (**g**) Baseline subtracted FLAG signal plotted against the fraction of the line integral of HA intensity for the ‘3d 4-OHT’, ‘4d 4-OHT’, ‘5d 4-OHT’, ‘6d 4-OHT’ groups in **e**. Thick centerline, mean; darker boundary in the close vicinity of the thick centerline, standard error of mean; lighter boundary, standard deviation. (**h**) An example line plot of the FLAG signal plotted against the fraction of line integral of HA intensity (from the ‘5d 4-OHT’ group in **e**), showing the quantification of the fraction of line integral of HA intensity when FLAG signal begins to rise (blue dot). Gold dashed line, the FLAG signal at the center of XRI (as baseline); green dashed line, a line fitted to the initial rising phase of the FLAG signal (defined as the portion of FLAG signal between 10% to 50% of the peak FLAG signal); blue dot, intersection of the two dashed lines. (**i-l**) Scatter plots of the fraction of the line integral of HA intensity when the FLAG signal begins to rise versus the length of the XRI (**i**), the thickness of the XRI (**j**), the curvature of the XRI (**k**), and the ratio of the FLAG signal at the end to the FLAG signal at the center (**l**), for XRIs in the ‘5d 4-OHT’ group (the ‘5d 4-OHT’ group was randomly chosen for this analysis). Gray line, line fit from linear regression. (**m**) Fraction of line integral of HA intensity when FLAG signal begins to rise plotted against the time of 4-OHT treatment after gene delivery, for XRIs in **g**. The line integral of HA intensity was normalized to ‘1’ for day 7, the time of cell fixation and thus the end of XRI growth. Middle line in box plot, median; box boundary, interquartile range; whiskers, 10-90th percentile; black dot, mean; black line, linear interpolation of the means. *, P < 0.05; **, P < 0.01; ***, P < 0.001; Kruskal-Wallis analysis of variance followed by post-hoc Dunn’s test.

Next, we quantified the relationship between the times of 4-OHT treatment and the resulting FLAG immunofluorescence patterns on XRI assemblies in neurons. Because the XRI growth is bidirectional over the 7-day experiment, we defined the fractional cumulative HA expression (i.e., the normalized, unidirectional line integral of HA immunofluorescence starting outwards from the center of the XRI) at the center of the XRI as ‘0’ and at the end of the XRI as ‘1’ (see **Supplementary Figure 2** for details of quantification). We hypothesized that this measure, the fractional cumulative HA expression, would correspond to a calibratable measure of time, postulating HA-bearing monomers to be added to the protein chain at a rate independent of the presence of non-HA-bearing monomers (i.e., FLAG-bearing monomers here), at least over the time scale of this experiment. That is, when FLAG-bearing monomers are being created, HA-bearing monomers are still being added to the growing protein chain at their own rate, although they are landing at more distant places along the chain, because FLAG-bearing monomers that were already added to the chain would lengthen the distance at which new HA-bearing monomers would land. Is this a reasonable postulate? We did see HA intensity to significantly decrease towards the end of XRI, when FLAG intensity increased due to 4-OHT induced expression of FLAG-bearing monomers (see ‘3-6d 4-OHT’ groups the first row in **Figure 2e**). In addition, this decrease in HA intensity towards the end of XRI was not observed without 4-OHT treatment (see ‘No-4-OHT’ group in the first row in **Figure 2e**). Because the 1POK-mediated fiber assembly has a fixed longitudinal monomer-to-monomer distance (~4 nm from electron microscopy measurements)^17^, the above results suggest that FLAG-bearing monomers took over a significant amount of longitudinal space at the end of XRI and thus diluted the line density of HA-bearing monomers.

This raises the question: is the assumption that HA-bearing and FLAG-bearing monomers are adding independently, each at a rate independent of the presence of the other monomer, a good one? If the binding and retention of HA-bearing monomers and FLAG-bearing monomers onto the XRI are both rare enough in time, that the chance of both types of monomers competing for the same slot on the XRI is insignificant, then this would be plausible. And, in this case, the fractional cumulative HA expression would still be a proper, calibratable measure of time. That is, if units with a new tag are supplementing the units being constitutively synthesized bearing an old tag, the latter units would not be added at a slower rate (i.e., there is no competition between the new units and the old units for being added to the growing chain), but instead would be added at the same rate, but simply be spaced out further from each other, separated by the units bearing the new tag. This would make the line integral the appropriate measure for extracting absolute time measurements. We sought to empirically test the hypothesis that absolute time measurements could be extracted from this specific measure. We averaged the FLAG signals across the two halves of the XRI (since XRIs are symmetric), to obtain the final FLAG signal (**Figure 2e,** bottom). Then, we calculated the ratio of the FLAG signal at the end of the XRI to the FLAG signal at the center of the XRI (**Figure 2f**), confirming that the polarized patterns of FLAG immunofluorescence on XRIs are present in neurons with 4-OHT treatments 3, 4, 5, or 6 days after AAV transduction, but not in neurons with 4-OHT treatments 1 or 2 days after AAV transduction. Therefore, we further analyzed the XRIs in neurons with 4-OHT treatments 3, 4, 5, or 6 days after AAV transduction, to characterize the relationship between the time of 4-OHT treatment and the fraction of the line integral of HA intensity at which the FLAG signal began to rise.

To quantify the fraction of the line integral of HA intensity at which the FLAG signal began to rise, we generated the net waveform of the FLAG signal with respect to the fraction of the line integral of HA intensity, by subtracting the baseline (i.e., the FLAG signal when the fraction of the line integral of HA intensity is zero) from the FLAG signal (**Figure 2g**). Next, we extrapolated the initial rising phase of the FLAG signal (defined as the period over which the FLAG signal increased from 10% to 50% of its peak value) until it intersected the pre-rising phase baseline (**Figure 2h**). The fraction of the HA line integral at this intersection point was defined as the point in time (although of course, to pinpoint a numerical value for the time requires calibration, discussed below) at which the FLAG signal began to rise. Importantly, this point did not depend on the length, thickness, or curvature of the XRI (**Figure 2i-k**), nor did it change with the precise value of the ratio of the FLAG signal at the end of the XRI to the FLAG signal at the center of the XRI (**Figure 2l**) – implying that this measure of time was a robust measure, and not dependent on the details of the geometry of the cell, and any associated constraints on the formation of the XRI. We also did not observe any correlation between the length, thickness, and curvature of XRI (**Supplementary Figure 3**), implying a certain degree of robustness as to the independence of different XRI geometrical attributes. As the time of 4-OHT treatment time increased, the fraction of the line integral of HA intensity when the FLAG signal began to rise also increased, albeit at a non-constant (i.e., increasing) rate, suggesting that the expression rate of AAV delivered XRI genes increased over time (**Figure 2m**). Thus time of a given cellular event can indeed be extracted from XRI geometry and label density, analyzed thus. We normalized this value to be 1 on day 7, because that was the time of cell fixation and thus the end of XRI growth (see day 7 in **Figure 2m**). We also replicated this experiment and applied expansion microscopy^23^ (ExM; a super-resolution imaging technique) instead of confocal microscopy for immunofluorescence imaging of XRI, obtaining similar results (**Supplementary Figure 4**). Thus, the predictable relationship between time of drug administration, and the fraction of the line integral of the HA intensity at which the FLAG signal began to increase, enables us to calibrate time information in XRI data analysis.

We next explored if XRIs could be used to record gene expression timecourse under mammalian immediate early gene (IEG) promoter activation. IEG promoters, such as the c-fos promoter^24^, are widely used to couple the expression of reporter proteins to specific cellular stimuli^25^. By using the c-fos promoter to drive the expression of XRI subunits tagged by a unique epitope tag, here the V5 tag, the time course of c-fos promoter driven expression could be recorded along the XRI filament, and read out by measuring the intensity profiles of V5 immunostaining signals along the filament. We chose to use the V5 tag here, instead of the previously used FLAG tag, so that each new XRI construct would be tagged by a unique epitope tag: in future usage of XRIs, one may want to co-express multiple XRI constructs in the same cell to achieve multiplexed recording of several different kinds of biological signals, readable via multiplexed immunostaining against distinct epitope tags. We expressed HA-bearing XRI, driven by the UBC promoter, in neurons using AAV as in the experiments in **Figure 2**, along with the new V5-bearing XRI driven by the c-fos promoter (**Figure 3a-c**). We diluted the AAV for the V5-bearing XRI (the final titer was 25% of that of the AAV for the HA-bearing XRI) so that the expression of HA-bearing monomers (and thus the HA portion of the final XRI assembly) would dominate over V5-bearing ones, and serve as a reliable integral substrate. We stimulated the neurons with 55 mM KCl, a common method to induce neuronal depolarization, for 3 hours, known to result in an increase in c-fos expression^26–28^. As expected, in the KCl stimulated neurons we observed low V5 immunofluorescence at the middle of the XRI, and towards both ends of the XRI the V5 immunofluorescence increased, resulting in peak-like patterns on each of the two sides of the XRI, eventually falling off (**Figure 3d,e**, right). This peak-like pattern of V5 immunofluorescence was not observed in XRIs in neurons without KCl stimulation (**Figure 3d,e**, left). The HA intensity fluctuated the opposite way of the V5 intensity (**Figure 3d, 3e**, right), as expected because, as discussed earlier, V5-bearing monomers would dilute down the line density of HA-bearing monomers; as long as the new V5 units being added were not competing with HA units being added, but simply were spacing the HA units out further, the line integral of HA units being added would be a useful measure of absolute time, at least over the time scale of this experiment (see above). Using the relationship between time and line integral of HA intensity obtained above (**Figure 2m**), we plotted relative change of V5 signal from baseline (baseline defined as the V5 signal when the fraction of the line integral of HA intensity was zero) along the XRI versus time. As expected, a peak of V5 signal was observed after the recovered time of day 5, which matched the actual time of KCl stimulation (**Figure 3e**, bottom row), while in neurons without KCl stimulation the V5 signal stayed relatively unchanged, a significant difference between these two cases (**Figure 3f**).

**Figure 3.**
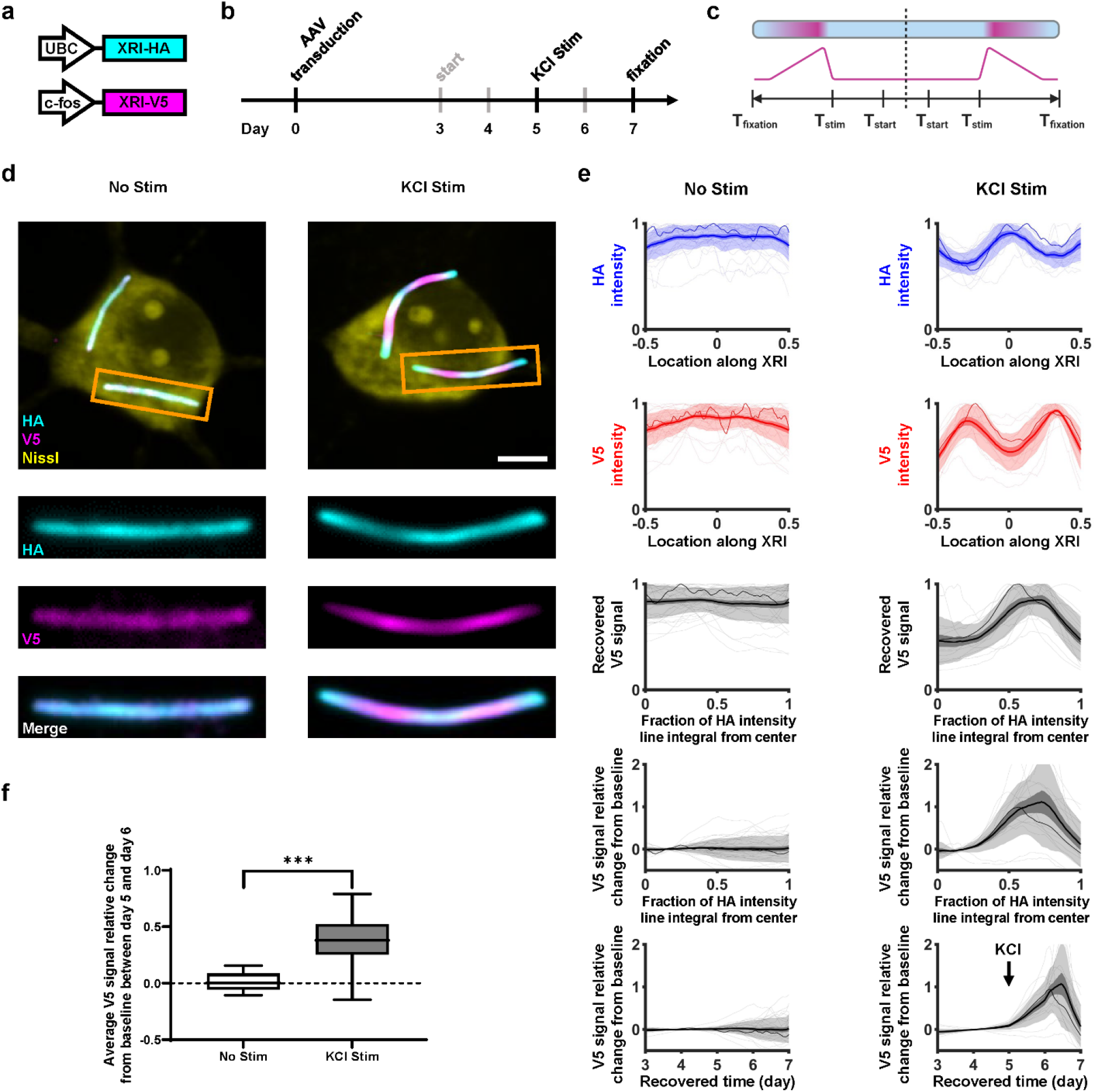
Recording the time course of c-fos promoter-driven expression with XRI. (**a-c**) Schematics of the AAV constructs co-transduced to neurons (**a**), experiment pipeline (**b**), and expected epitope distribution along the XRI protein self-assembly (**c**) in the c-fos promoter-driven gene expression experiment. XRI-HA, XRI with the epitope tag HA; XRI-V5, XRI with the epitope tag V5; c-fos, c-fos promoter; T_stim_, the time of the onset of stimulation of neuron activity by KCl; T_start_, the time when XRI starts recording information after gene delivery and expression of XRI, which is measured to be 3 days after AAV transduction in **Figure 2**. (**d**) Representative confocal images of cultured mouse hippocampal neurons expressing constructs in **a**, taken after fixation, Nissl staining, and immunostaining against HA and V5 tags. KCl stim, 55 mM KCl stimulation for 3 hours starting at T_stim_ = 5 days; three rows of rectangular panels at the bottom, enlarged views of regions marked in orange rectangles in the top row of square panels. Scale bar, 5 μm. (**e**) HA intensity profile along the XRI (first row), V5 intensity profile along the XRI (second row), recovered FLAG signal (by averaging the two FLAG intensity profiles across the two halves of the XRI) plotted against the fraction of the line integral of HA intensity (third row), V5 signal relative change from baseline (ratio of the V5 signal to the V5 signal at the center of the XRI) plotted against the fraction of the line integral of HA intensity (fourth row), and V5 signal relative change from baseline plotted against recovered time (using the black line in **Figure 2m** as time calibration for time recovery from the line integral of HA intensity; fifth row), from the experiment described in **a-c** (n = 30 XRIs from 28 neurons from 2 cultures for ‘No Stim’ group; n = 15 XRIs from 11 neurons from 2 cultures for ‘KCl Stim’ group). Thick centerline, mean; darker boundary in the close vicinity of the thick centerline, standard error of mean; lighter boundary, standard deviation; lighter thin lines, data from individual XRIs; darker thin line, data from the corresponding XRI in the orange rectangle in **d**. In the first three rows, each raw trace was normalized to its peak to show relative changes before averaging. See **Supplementary Figure 2** for the detailed process flow of extracting signals from XRI assemblies. (**f**) Box plot of the average V5 signal relative change from baseline between day 5 and day 6 (i.e., within 24 hours after the onset time of KCl stimulation). Middle line in box plot, median; box boundary, interquartile range; whiskers, 10-90 percentile; black dot, mean. ***, P < 0.001; Wilcoxon rank sum test. See **Supplementary Table 3** for details of statistical analysis.

Next, we tested if XRI can preserve temporal information in the living mammalian brain. We took the same XRI AAVs used in **Figure 2** and co-injected them into the hippocampal CA1 region of the brains of adult wild-type mice (**Figure 4a,b**). Based on previous experience from us and others^29,30^ on the AAV-mediated gene delivery of Cre (in the experiment here ERT2-iCre-ERT2 was delivered) into the mouse brain *in vivo*, we doubled the expression time to 14 days for this *in vivo* experiment, so that 4-OHT was administered into mouse via intraperitoneal injection^31^ at 10 days after AAV injection (5/7 of the way through the experimental timecourse) to induce the enzymatic activity of ERT2-iCre-ERT2, which triggers the expression of the FLAG-bearing XRI, and then the mouse brain was fixed and sectioned 14 days after AAV injection for downstream immunofluorescence (see the experiment pipeline in **Figure 4b**). After immunofluorescence imaging of the resulted brain slices, we observed abundant expression of XRI in neurons in the CA1 area (**Figure 4c**; see images of individual representative neurons in **Figure 4d**). Similar to what was observed in cultured neurons in **Figure 2**, the FLAG immunofluorescence had a strong polarized pattern in the XRIs formed *in vivo*, confirming that XRI can indeed preserve temporal information in the living mammalian brain.

**Figure 4.**
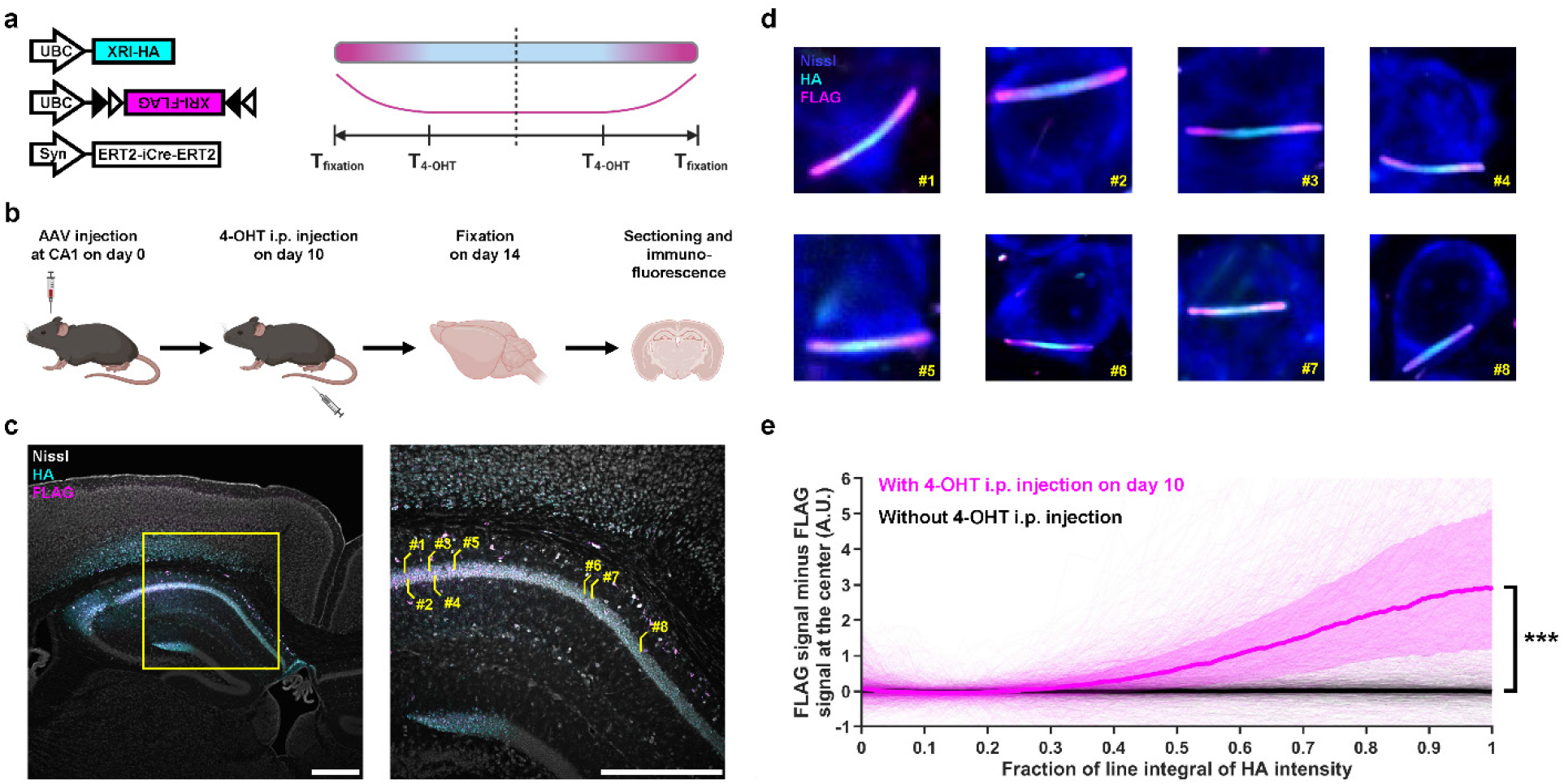
*In vivo* XRI self-assembly in mouse brain. (**a-b**) Schematics of the AAV constructs (**a**, left), expected epitope distribution along the XRI protein self-assembly (**a**, right), and experiment pipeline (**b**) in this XRI self-assembly experiment in mouse brain. AAVs were injected into the dorsal CA1 area of the brains of 3-month-old mice on day 0, followed by 4-OHT intraperitoneal (i.p.) injection on day 10 and then fixation via 4% paraformaldehyde perfusion on day 14. The preserved brains were then sectioned at 50 μm coronally and stained with anti-HA, anti-FLAG, and Nissl stain. (**c**) Confocal images of a representative brain section from the experiment described in **b**. Yellow square on the left panel, boundary of the region of interest enlarged in the right panel; lines and numbers on the right panel, locations of the neurons shown in **d**; scale bars, 500 μm. (**d**) Confocal images of representative CA1 neurons indicated in the right panel in **c**. (**e**) FLAG signal minus the FLAG signal at the center averaged and plotted against the fraction of the line integral of HA intensity along the XRI. n = 893 XRIs from 835 CA1 neurons from 1 brain section (the one shown in **c**) from 1 mouse with 4-OHT i.p. injection on day 10 (shown in magenta) and n = 598 XRIs from 475 CA1 neurons from 1 brain section from 1 mouse without 4-OHT i.p. injection (shown in black). The line integral of HA intensity was defined as ‘1’ for day 14, the time of fixation and thus the end of XRI growth. Colored lines, median; colored, shaded boundaries, interquartile range; lighter thin lines, data from individual XRIs. ***, P < 0.001; Wilcoxon rank sum test. See **Supplementary Table 3** for details of statistical analysis.

We analyzed the XRIs in 835 CA1 neurons in confocal imaged volumes and plotted the absolute, baseline subtracted (baseline defined as the signal at the center of XRI) FLAG signals with respect to the fraction of the line integral of HA intensity, and performed the same analysis on XRIs in 475 CA1 neurons in another mouse that underwent the same experimental pipeline but without 4-OHT injection (**Figure 4e**). FLAG signals in the mouse without 4-OHT injection were flat with respect to the fraction of the line integral of HA intensity, while those in the mouse with 4-OHT injection on day 10 began to rise when the fraction of the line integral of HA intensity reached 0.3. This 0.3 value alone does not provide absolute information about the time axis, without an *in vivo* calibration of the timecourse as done *in vitro* for **Figure 2** – but, we note that that this 0.3 value, from this day 10 4-OHT injection amidst a 14-day *in vivo* experiment, matched the same value obtained for the day 5 4-OHT treatment in the 7-day experiment in cultured neurons (**Figure 2m**). Note that in both cases, 4-OHT was given at a time point 5/7 of the way through the total XRI expression time, suggesting that this time point corresponds to 30% of the fraction of the line integral of HA intensity, in multiple neural preparations. Future work on developing XRI for *in vivo* use should replicate the calibration experiment of **Figure 2m** in the living mouse brain, to precisely numerically calibrate the time axis.

## Discussion

In this work, we proposed and experimentally confirmed that cellular physiological information could be recorded onto intracellular, steadily growing, protein chains made out of fully genetically encoded self-assembling proteins, and then read out via routine immunofluorescence and imaging techniques. By screening existing, human-created self-assembling protein candidates, and then performing protein engineering to add an “insulator” component to the promising self-assembling protein candidate 1POK to encourage stable, time-ordered longitudinal growth, we developed what we call an expression recording island (XRI), named by analogy to our earlier signaling reporter island technology (SiRI, which also uses self-assembling peptides, but in that case to create a spatial encoding of indicator identity^32^) -- a fully genetically encoded system for recording biological information via self-assembling protein chains. We defined, provided rationale for, and validated, a calibratable measure of time, the fractional cumulative expression of HA-bearing monomers, to calibrate the time axis onto the information recorded on the XRI via ordered epitope tags. We applied XRIs to record c-fos promoter-driven gene expression in cultured mouse hippocampal neurons after depolarization, and applied the fractional cumulative expression of HA-bearing monomers to recover the time axis and c-fos promoter-driven gene expression solely from information read out from XRI via immunostaining and imaging. We showed that XRI could preserve the temporal order of protein monomers expressed in the living mouse brain. Thus, XRIs function in multiple biological systems, including the live mammalian brain, in encoding cellular physiological signals into a linear, optically readable protein chain.

Compared to nucleic acid-based systems which require nucleic acid sequencing methods that require dissociation and/or lysis of cells^7–13^, reading out recorded information from a protein-based system through imaging only requires routine immunofluorescence techniques and conventional microscopes, available to many biology groups already, without the need for additional hardware investment. Such preservation of cellular physiological information within the native environment offered by our protein-based system also would enable correlation of the recorded biological information with other kinds of structural and molecular information associated with the cellular population, such as the spatial location, cell type, and presence of protein and other markers in the recorded cells^5,6,33^, some of which may be causally involved with the creation of the physiological signals, or that result from the physiological signals. Such kinds of multimodal data could enable the analysis of how specific cellular machinery drive, or result from, complex timecourses of physiological stimuli. For example, by offering the ability to record gene expression time course in single cells, as shown here, the proposed protein-based XRI system will enable the study of gene expression time course as a result of specific cellular inputs and/or drug treatments^34,35^. This could be useful, amongst many other possibilities, for the investigation of circadian gene rhythms^36^ and rhythms of other genes that change in complex ways over time. XRIs could be used to record transcription factor activities^37^, or as an information storage platform to externally introduce unique cellular barcodes into single cells for cell identification^38^, as just a few out of many possibilities.

Future work may include the development of mechanisms for coupling XRI expression to other biological dynamics and processes, which would significantly broaden the kinds of biological information XRI could record. For example, the c-fos promoter we used in the study is a natural “tool” that couples c-fos promoter activity to XRI expression. Ongoing activities to engineer promoters and expression systems that respond to calcium^39,40^ and other physiological dynamics^25,41^, or stages of the cell cycle^42^, would enable XRI recordings of calcium activity or cell cycle. Another future direction will be to expand the XRI system for multiplexed recording of multiple kinds of biological information onto the same polymer chain, using a unique epitope tag for each kind of biological information, and multiplexed immunostaining methods^32^ to read out each information. For example, one could use tags A and B to encode the gene expression history of genes 1 and 2, respectively, and use tag C to encode the calcium signal, by expressing all the components simultaneously, and then immunostaining all the tags after fixation. Future work may also improve XRI designs to reach time resolutions of recording well below ~1 day, perhaps even towards minute timescales or better.

---

**Supplementary Figure 1.**
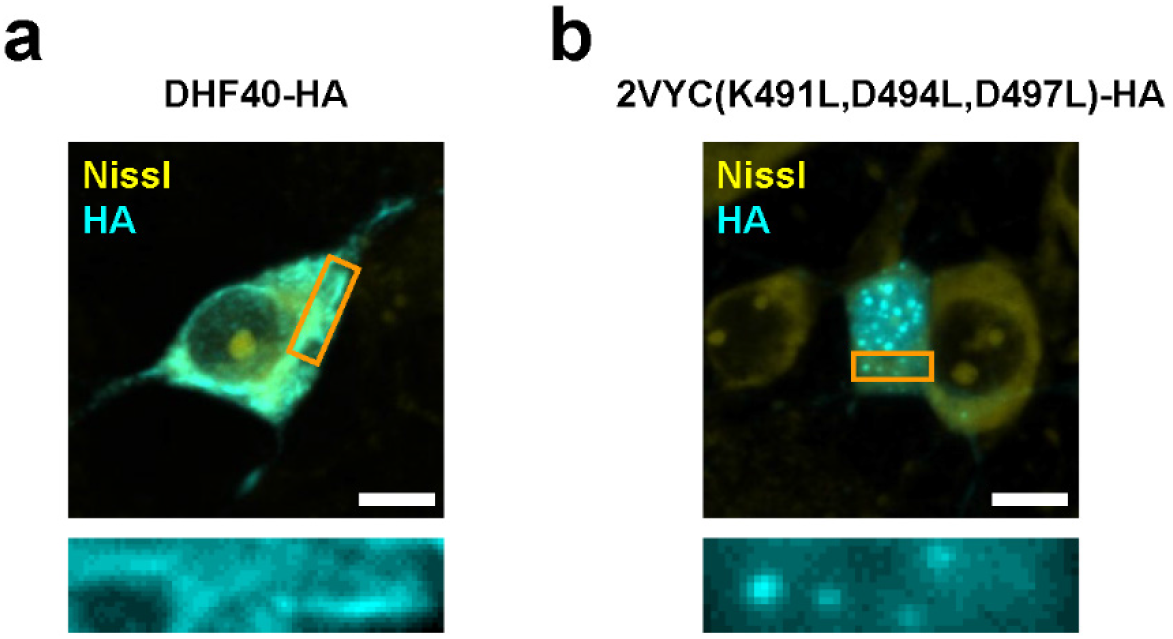
Testing additional self-assembling protein constructs in neurons. Representative confocal images of cultured mouse hippocampal neurons expressing (**a**) DHF40 or (**b**) 2VYC(K491L,D494L,D497L) with epitope tag HA, taken after fixation, Nissl staining, and immunostaining against the HA tag. Scale bars, 5 μm throughout this figure. Rectangular panels at the bottom, enlarged views of regions marked in orange rectangles in the top row of square panels. See **Supplementary Table 1** for sequences of the motifs; see **Supplementary Table 2** for all tested constructs.

**Supplementary Figure 2.**
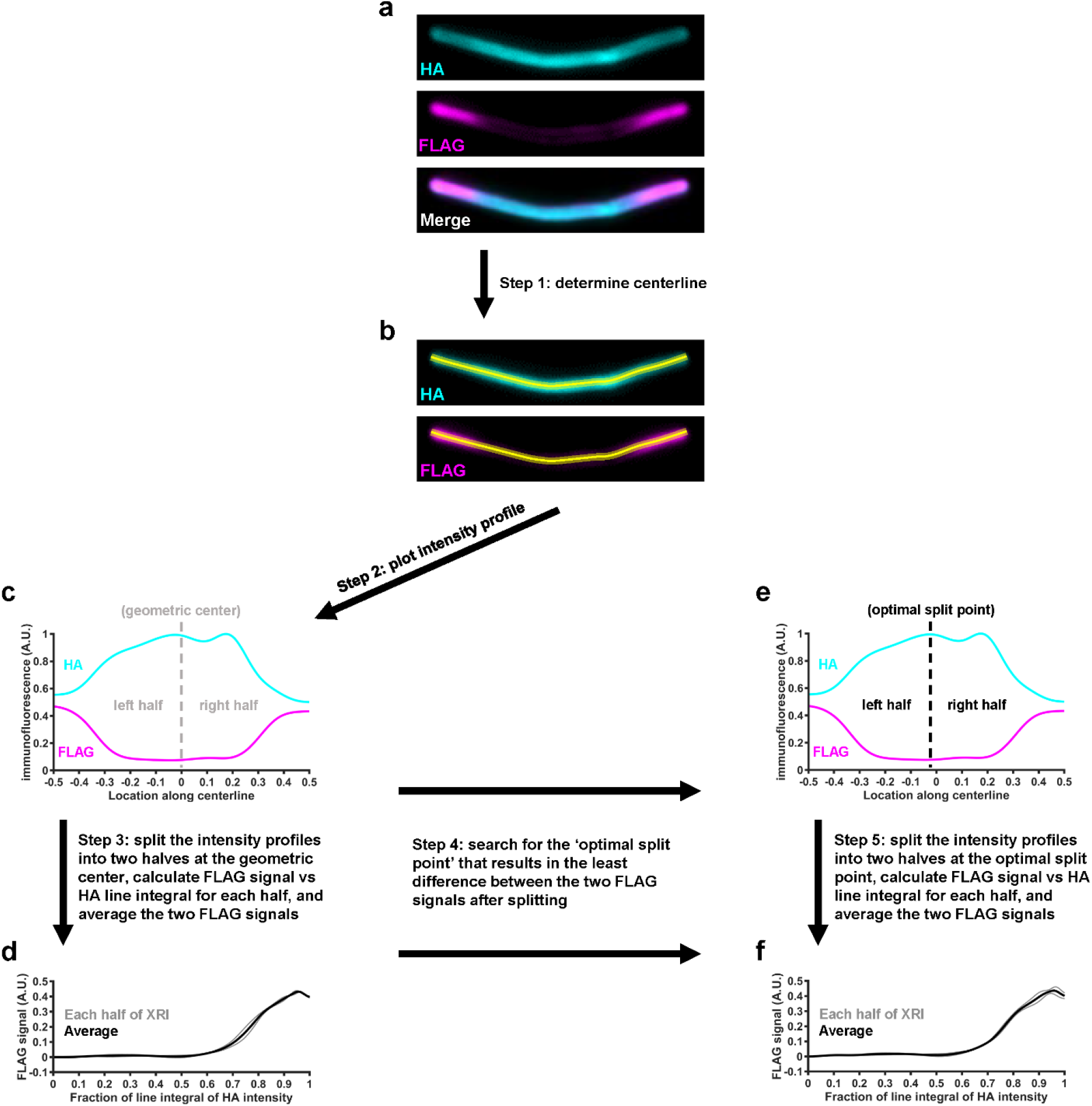
Process flow for extracting information from XRI assemblies. **Step 1:** For each XRI (**a**), a curved centerline was drawn along the longitudinal axis of the XRI in the anti-HA channel (**b**). The centerline width was set to half of the width of the XRI. **Step 2:** The intensity profiles along this centerline were measured in the anti-HA channel (resulting in an HA line profile; cyan curve in **c**) and in the other XRI epitope staining channel, such as in the anti-FLAG channel (resulting in a FLAG line profile; magenta curve in **c**). **Step 3:** Next, each of the line profiles was split into two half line profiles using the geometric center point of the XRI (the 50% length point along the centerline, measuring from the end of the XRI; gray dashed vertical line in **c**) as the ‘split point’. Each of the half HA line profiles was then converted into a line integral of HA, by integrating the line profile with respect to the distance along the half centerline starting from the geometric center point, and then these line integrals of HA were normalized to the maximum integral value so that each line integral of HA started at the value 0 at the geometric center point of the XRI, and gradually increased to the value 1 at the end of the XRI. For the corresponding half FLAG line profiles, line integrals were also calculated but not normalized. At this point, we have the line integrals of HA and FLAG, which correspond to the cumulative HA and FLAG intensities along each half of the XRI. The FLAG intensity change per unit change in the cumulative HA intensity, defined as the FLAG signal, was calculated by taking the derivative of the line integral of FLAG with respect to the line integral of HA (gray curves in **d**). At this stage, we had obtained the line integral of HA and the FLAG signal from each of the halves of the XRI, and the final extracted FLAG signal from this XRI (black curves in **d**) was defined as the point-by-point average of the two FLAG signals from the two halves of the XRI. **Step 4:** We found the two obtained FLAG signals from the same XRI have small but noticeable differences (see the two gray curves in **d**). We reasoned that such small but noticeable discrepancies between the two halves of the same XRI was due to the asymmetry of the XRI, and the choice of the exact geometric center as the split point may not be optimal. To minimize the discrepancy between the two FLAG signals from the two halves of the same XRI, we searched for an optimal split point (black dashed vertical line in **e**) near the geometric center of the XRI (searching range was the geometric center +/- 10% of the total XRI length, i.e., between −0.1 and 0.1 on the x-axis in **e**), so that using this optimal split point, instead of the geometric center, as the split point would result in the least difference (in terms of sum of squared differences) between the two FLAG signals from the two halves of the split XRI. **Step 5:** Same as **Step 3**, except that the optimal split point, instead of the geometric center, was used to split the intensity profiles into two halves (**f**). We found the resulting final FLAG signal (after averaging those from the two halves) when using the geometric center as the split point was similar to that when using the optimal split point as the split point (compare the black line in **d** and **f**). Nevertheless, we used the optimal split point as the split point to analyze XRIs throughout this paper.

**Supplementary Figure 3.**
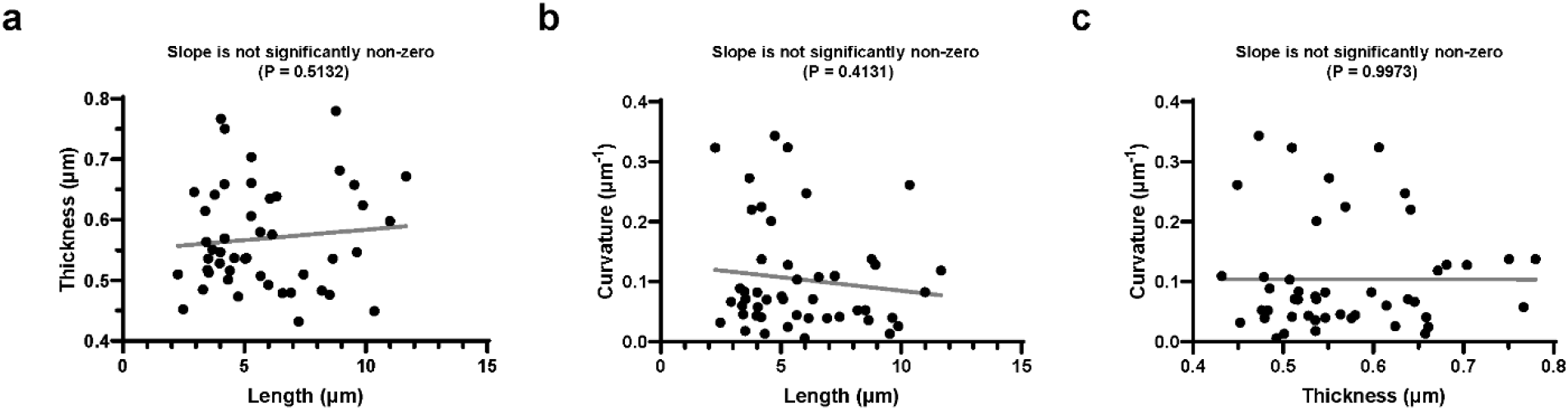
Additional geometric analysis of XRI. (**a**) Scatter plot of the thickness of XRI versus the length of XRI (in neurons in the ‘5d 4-OHT’ group in **Figure 2** throughout this figure; n = 47 XRIs from 32 neurons from 2 cultures). Gray line, line fit from linear regression throughout this figure. (**b**) Scatter plot of the curvature of XRI versus the length of XRI. (**c**) Scatter plot of the curvature of XRI versus the thickness of XRI.

**Supplementary Figure 4.**
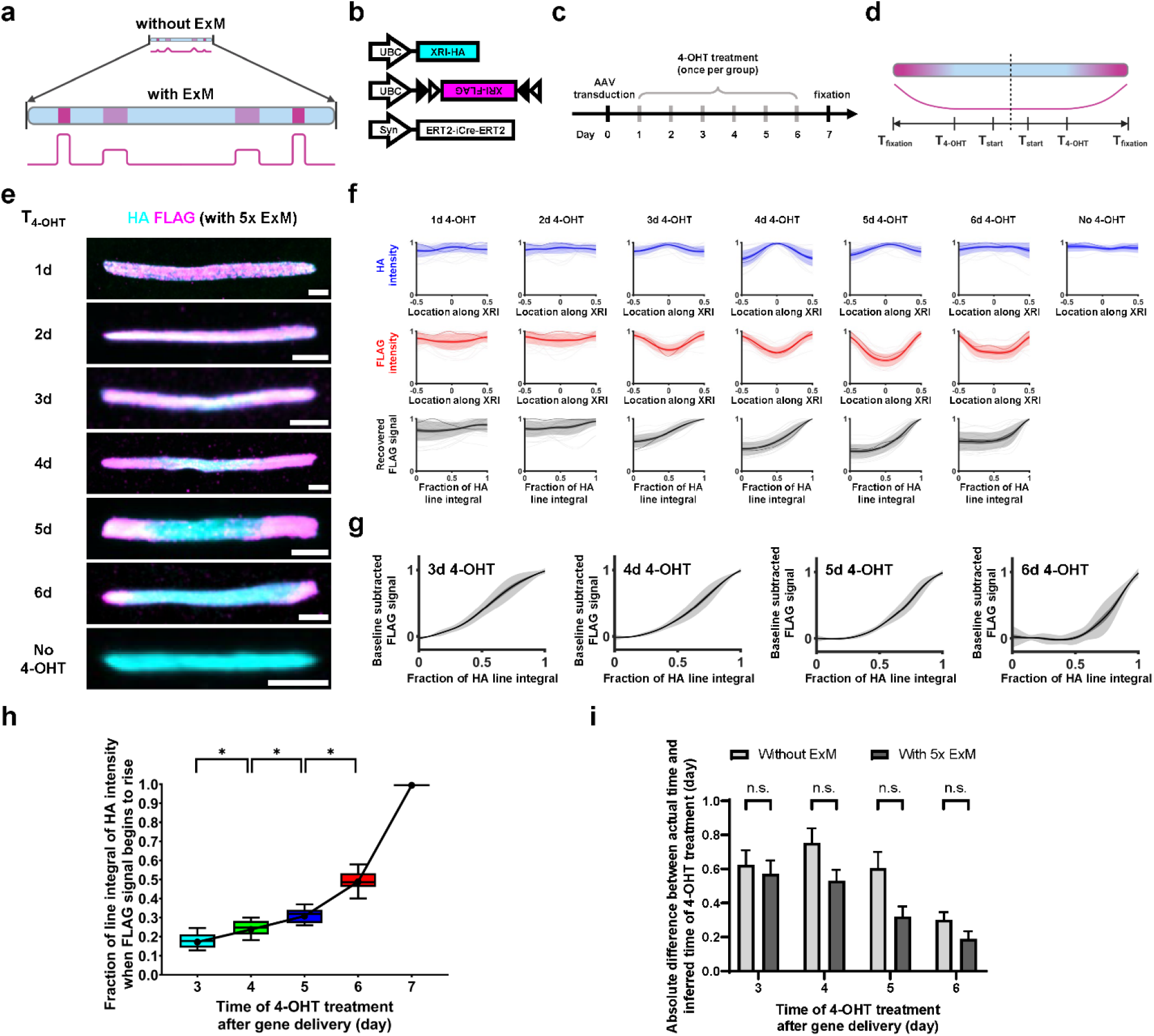
Imaging XRIs using expansion microscopy. We replicated the experiment in **Figure 2** and applied expansion microscopy^23^ (ExM) instead of confocal microscopy for immunofluorescence imaging of XRIs. We optimized the digestion methods (removing the proteinase K digestion step and replacing it with a heat-based softening step) starting from the TREx^43^ ExM protocol, while receiving inspirations from the ExR^44^ protocol, to achieve uniform expansion of XRI assemblies and post-expansion immunostaining/immunofluorescence at a high signal-to-noise-ratio (at a linear expansion factor of ~ 5x), with antibody staining against NeuN to locate the somata of neurons. (**a**) Schematic of using expansion microscopy (ExM) to increase the spatial resolution of immunofluorescence imaging. (**b-d**) Schematics of the constructs co-transduced to neurons (**b**), experiment pipeline (**c**), and expected epitope distribution along the XRI protein self-assembly (**d**) in the chemically induced gene expression experiment, as in **Figure 2**. (**e**) Representative confocal images of XRIs in cultured mouse hippocampal neurons expressing constructs in **b** with different times of 4-OHT treatment (T_4-OHT_), after 5x ExM. Scale bars, 5 μm after ExM (equivalent to 1 μm in biological units, e.g. when divided by the expansion factor). (**f**) HA intensity profile along the XRI (top row), FLAG intensity profile along the XRI (middle row), and recovered FLAG signal (by averaging the two FLAG intensity profiles across the two halves of XRI), plotted against the fraction of the line integral of HA intensity (a value between 0 and 1; 0 corresponds to the center of XRI, and 1 corresponds to the end of XRI; bottom row), from the experiment described in **a-d** (n = 32 XRIs from 19 neurons from 2 cultures for ‘1d 4-OHT’ group; n = 30 XRIs from 16 neurons from 2 cultures for ‘2d 4-OHT’ group; n = 23 XRIs from 14 neurons from 2 cultures for ‘3d 4-OHT’ group; n = 24 XRIs from 15 neurons from 3 cultures for ‘4d 4-OHT’ group; n = 22 XRIs from 17 neurons from 3 cultures for ‘5d 4-OHT’ group; n = 19 XRIs from 15 neurons from 3 cultures for ‘6d 4-OHT’ group; n = 7 XRIs from 3 neurons from 1 culture for ‘No 4-OHT’ group). Each raw trace was normalized to its peak, to show relative changes, before averaging. Thick centerline, mean; darker boundary in the close vicinity of the thick centerline, standard error of mean; lighter boundary, standard deviation; lighter thin lines, data from individual XRIs; darker thin line, data from the corresponding XRI in **e**. See **Supplementary Figure 2** for the detailed process flow of extracting signals from XRI assemblies. (**g**) Baseline subtracted FLAG signal plotted against the fraction of the line integral of HA intensity for the ‘3d 4-OHT’, ‘4d 4-OHT’, ‘5d 4-OHT’, ‘6d 4-OHT’ groups in f. Thick centerline, mean; darker boundary in the close vicinity of the thick centerline, standard error of mean; lighter boundary, standard deviation. (**h**) Fraction of line integral of HA intensity when FLAG signal begins to rise, plotted against the time of 4-OHT treatment after gene delivery, for XRIs in **g**. The line integral of HA intensity was normalized to ‘1’ for day 7, the time of cell fixation and thus the end of XRI growth. Middle line in box plot, median; box boundary, interquartile range; whiskers, 10-90 percentile; black dot, mean; black line, linear interpolation of the means. *, P < 0.05; Kruskal-Wallis analysis of variance followed by post-hoc Dunn’s test. See **Supplementary Table 3** for details of statistical analysis. (**i**) Bar plot of the absolute difference between the actual time and the inferred time of 4-OHT treatment, without and with 5x ExM. For each XRI, the inferred time of 4-OHT treatment was calculated from the fraction of the line integral of HA intensity when the FLAG signal begins to rise, using the black line in **Figure 2m** (for XRI without ExM) or the black line in **h** (for XRI with 5x ExM) as time calibration. Bar height, mean; error bars, standard error of mean. n.s., not significant; Bonferroni corrected Wilcoxon rank sum tests.

## Acknowledgments

We thank Fan Wang, Mark Bear, Amy Keating, George Church, and Keith Tyo for useful discussions. C.L. acknowledges the J. Douglas Tan Postdoctoral Fellowship. E.S.B. was supported by Lisa Yang, John Doerr, NIH 1R01MH123977, NIH R01DA029639, NIH R01MH122971, NIH RF1NS113287, NSF 1848029, NIH UF1NS107697, NIH 1R01DA045549, NIH 1R01MH114031, U. S. Army Research Laboratory and the U. S. Army Research Office under contract/grant number W911NF1510548, and HHMI.

## Author Contributions

C.L. and E.S.B. made high-level plans, interpreted the data and wrote the paper. C.L. performed protein design with the help from W.M.P., and performed screening experiments. C.L. and N.S. performed live cell experiments and immunofluorescence of cultured cells and brain slices. B.A. performed expansion microscopy experiments with the help from C.Z. M.S. and S.R. performed mice experiments. C.L., B.A. and O.T.C. analyzed the data.

## Data Availability

The data sets generated and analyzed in the current study will be deposited to Zenodo Archive.

## Code Availability

The code used to produce analysis and figures for the current study will be deposited to Zenodo Archive.

## Supplementary Tables

**Supplementary Table 1.**
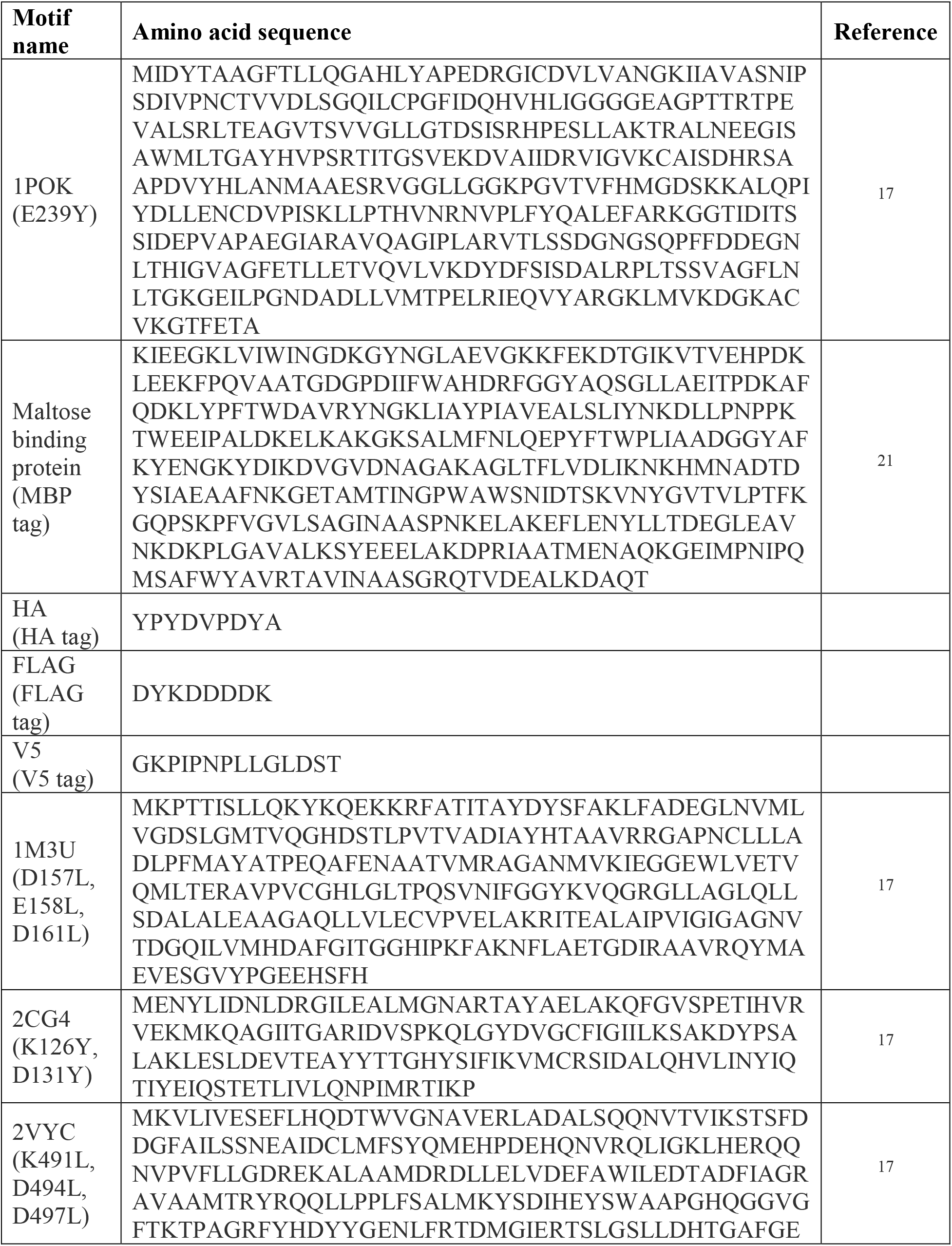
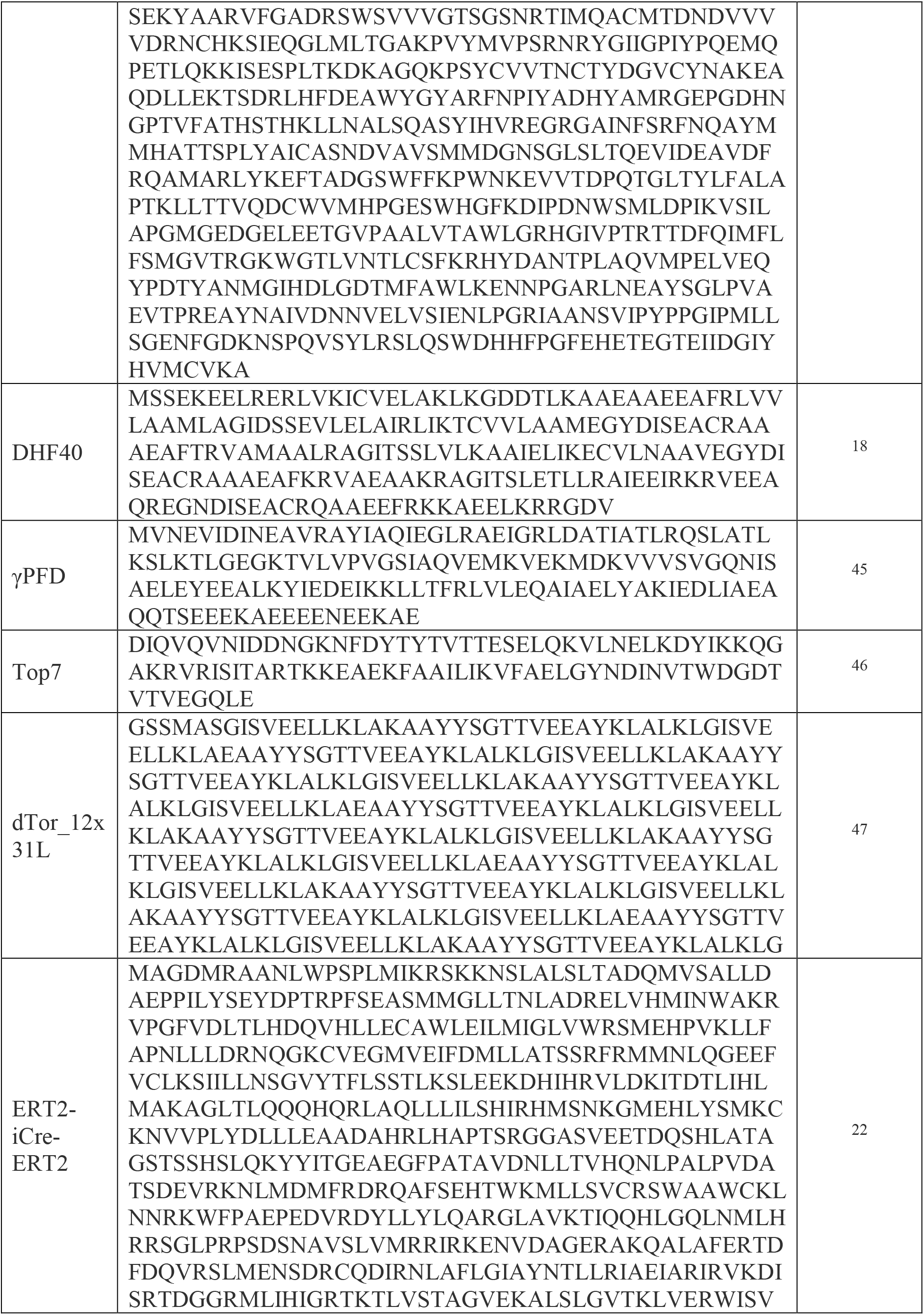
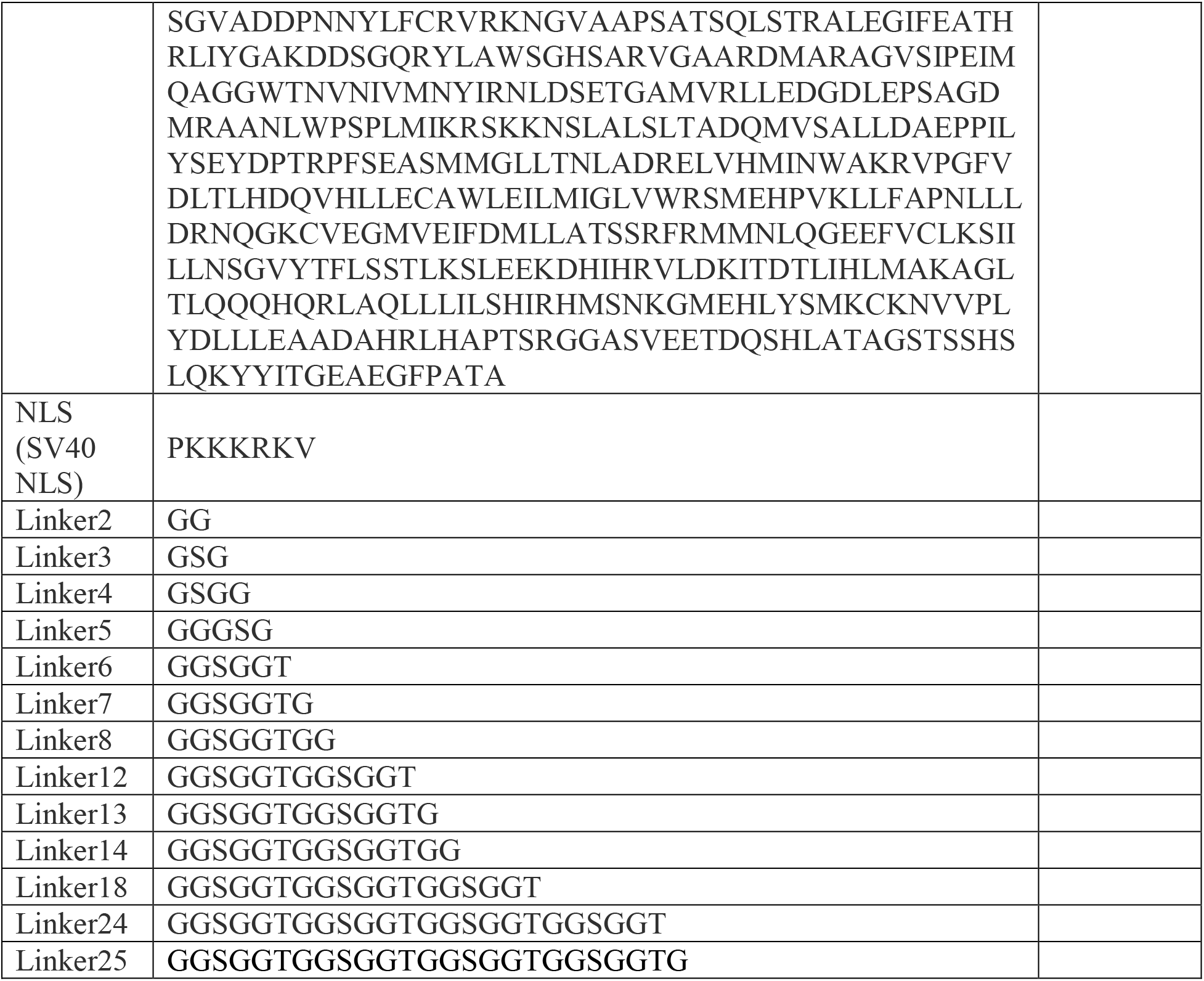
sequences of protein motifs used in this study

**Supplementary Table 2.** constructs of self-assembly proteins tested in neurons in this study

| **Construct** (promoters are <u>underlined</u>) | **Resulted pattern of protein self-assembly** (in the cytosol unless noted otherwise) |
| --- | --- |
| <u>UBC</u>-1POK(E239Y)-Linker25-HA-Linker3-MBP_tag<br>(also known as XRI-HA) | Fiber(s) |
| <u>UBC</u>-1POK(E239Y)-Linker12-gg-HA | Unstructured aggregates and intertwined fibers |
| <u>UBC</u>-1POK(E239Y)-Linker13-HA-mEGFP | Fiber(s) |
| <u>UBC</u>-1M3U(D157L,E158L,D161L)-Linker14-HA | Unstructured aggregates (and intertwined fibers in a subset of cells) |
| <u>UBC</u>-HA-Linker14-2CG4(K126Y,D131Y) | Uniform expression in the nucleus |
| <u>UBC</u>-2VYC(K491L,D494L,D497L)-Linker14-HA | Nucleus-localized puncta and cytosol-localized puncta |
| <u>CMV</u>-1POK(E239Y)-Linker8-HA | Unstructured aggregates and intertwined fibers |
| <u>UBC</u>-1POK(E239Y)-Linker7-HA-Linker3-MBP_tag | Fiber(s) |
| <u>UBC</u>-HA-Linker3-MBP_tag-Linker18-1POK(E239Y) | Fiber(s) |
| <u>UBC</u>-1POK(E239Y)-Linker5-HA-mEGFP | Fibers (mostly) and puncta |
| <u>UBC</u>-mEGFP-HA-Linker12-1POK(E239Y) | Fiber(s) |
| <u>UBC</u>-1POK(E239Y)-Linker25-HA-g-mEGFP | Fiber(s) |
| <u>UBC</u>-1POK(E239Y)-Linker7-HA-Linker3-Top7 | Short fibers and puncta in the nucleus (mostly) and cytosol |
| <u>UBC</u>-1POK(E239Y)-Linker25-HA-gsg-Top7 | Unstructured aggregates and intertwined fibers in the cytosol; nucleus-localized fibers |
| <u>UBC</u>-1POK(E239Y)-Linker5-mEGFP-Linker2-HA-Linker3-MBP_tag | Puncta |
| <u>UBC</u>-1POK(E239Y)-Linker24-mEGFP-HA-Linker6-MBP | Fiber(s) with large thickness |
| <u>UBC</u>-1POK(E239Y)-Linker5-mEGFP-HA-Linker3-Top7 | Dense and small puncta |
| <u>UBC</u>-Top7-Linker12-1POK(E239Y)-Linker13-HA-mEGFP | Unstructured aggregates and fibers; high non-assembly background |
| <u>UBC</u>-1POK(E239Y)-Linker24-mEGFP-HA-Linker6-Top7 | Unstructured aggregates and fibers in the cytosol; nucleus-localized fibers |
| <u>UBC</u>-HA-dTor_12x31L-Linker24-1POK(E239Y) | Puncta |
| <u>UBC</u>-NLS-Linker4-1POK(E239Y)-Linker13-HA-mEGFP | Nucleus-localized fiber(s) |
| <u>UBC</u>-NLS-Linker4-1POK(E239Y)-Linker14-HA | Nucleus-localized puncta |
| <u>UBC</u>-DHF40-Linker14-HA | Unstructured aggregates and intertwined fibers |
| <u>UBC</u>-DHF40-Linker13-HA-mEGFP | Unstructured aggregates, puncta, and intertwined fibers |
| <u>UBC</u>-DHF58Four-Linker14-HA | Unstructured aggregates (and intertwined fibers in a subset of cells) in the cytosol and nucleus |
| <u>UBC</u>-DHF58Six-Linker14-HA | Uniform expression in the nucleus, with dim unstructured aggregates in the cytosol |
| <u>UBC</u>-DHF58Six-Linker14-mRuby2_smFP(HA) | Uniform expression in the nucleus, with dim unstructured aggregates in the cytosol |
| <u>UBC</u>-DHF79-Linker14-HA | Uniform expression in the nucleus, with dim unstructured aggregates in the cytosol |
| <u>UBC</u>-DHF119-Linker14-HA | Uniform expression in the nucleus, with dim unstructured aggregates in the cytosol |
| <u>CMV</u>-DHF40-Linker8-HA | Unstructured aggregates and intertwined fibers |
| CMV-DHF46-Linker8-HA | Puncta |
| CMV-DHF47-Linker8-HA | Unstructured aggregates and puncta |
| CMV-DHF50-Linker8-HA | Unstructured aggregates and puncta |
| CMV-DHF77-Linker8-HA | Unstructured aggregates and puncta |
| UBC-γPFD-Linker8-HA | Puncta |

**Supplementary Table 3.**
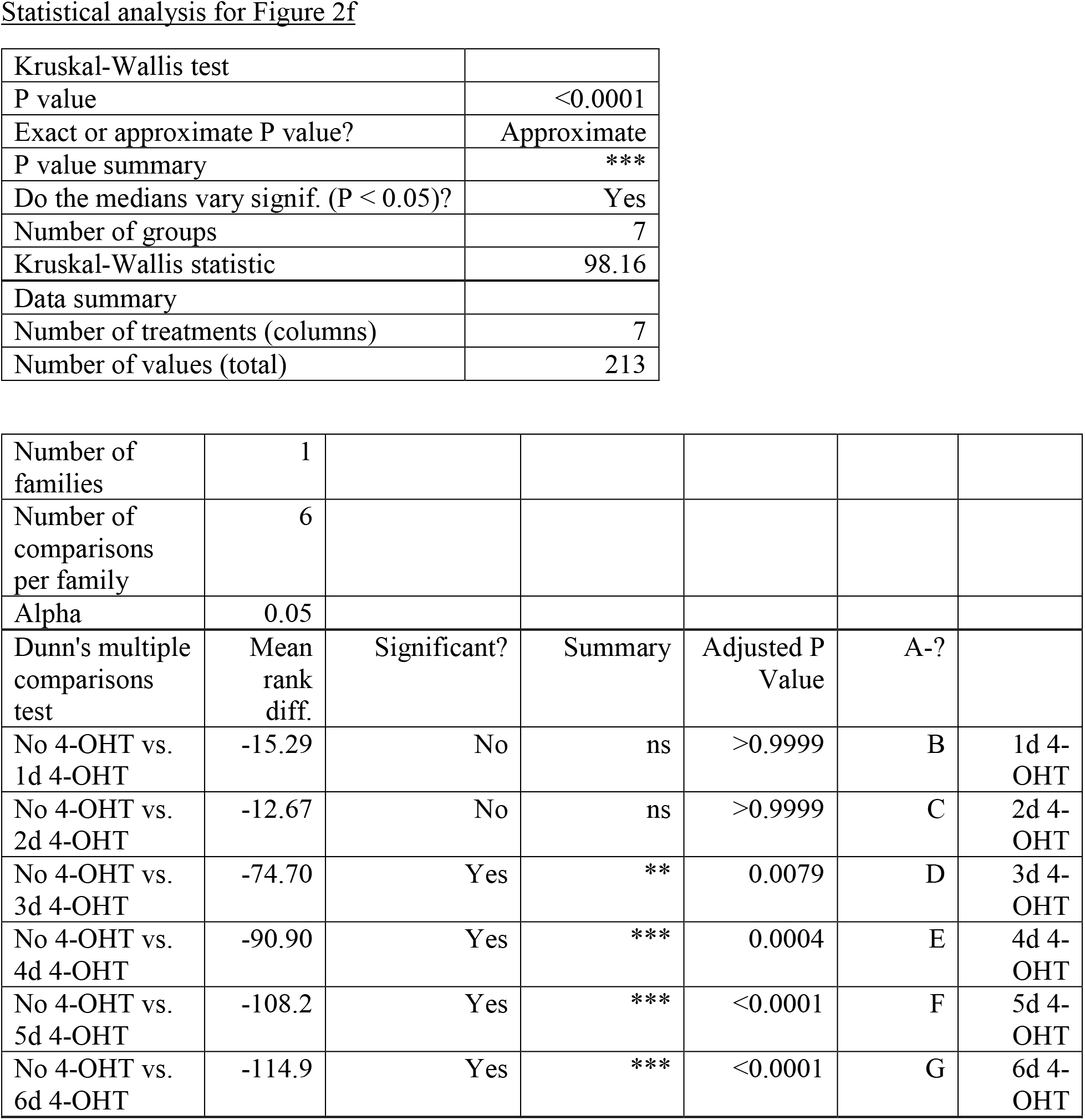
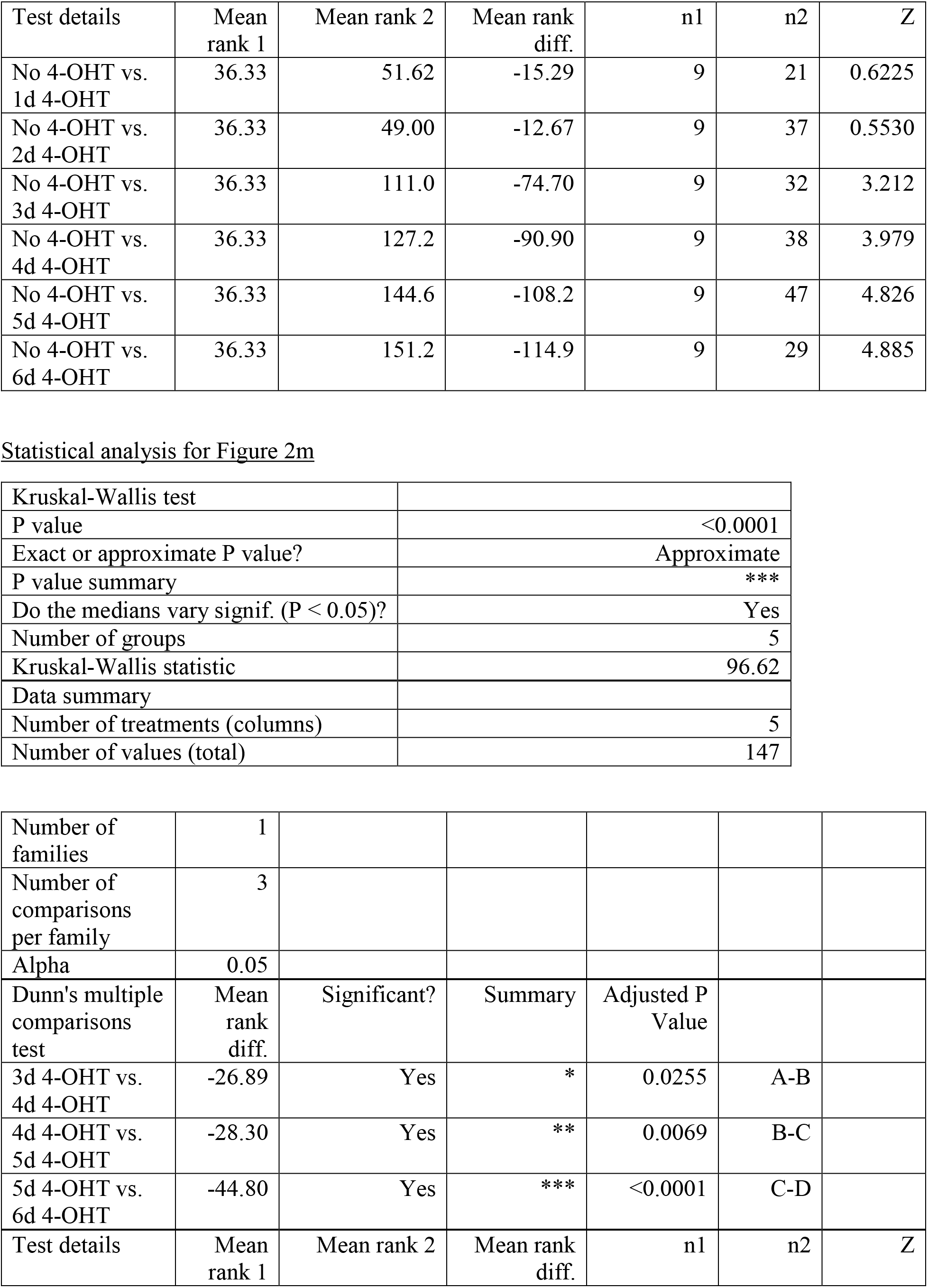
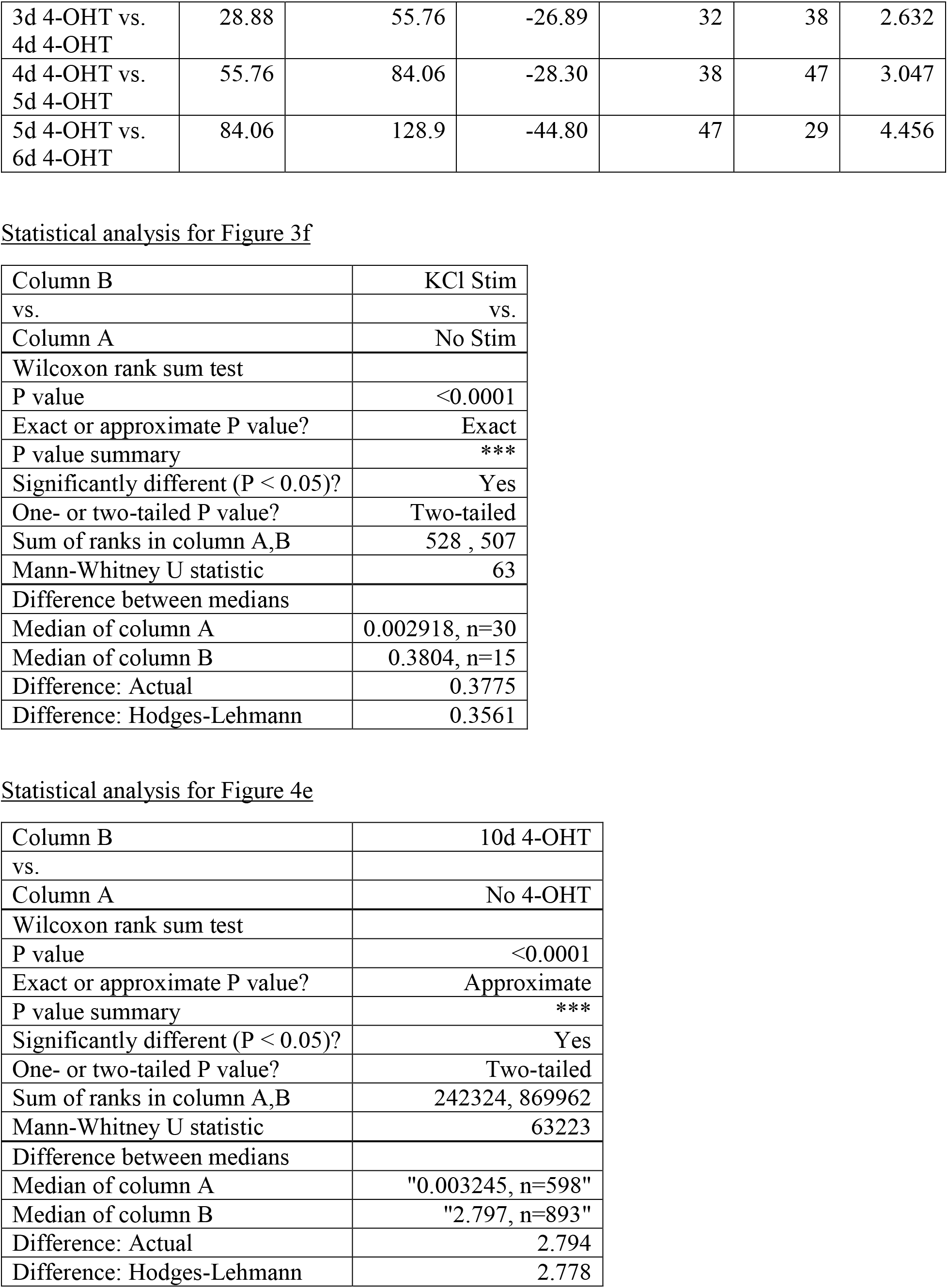
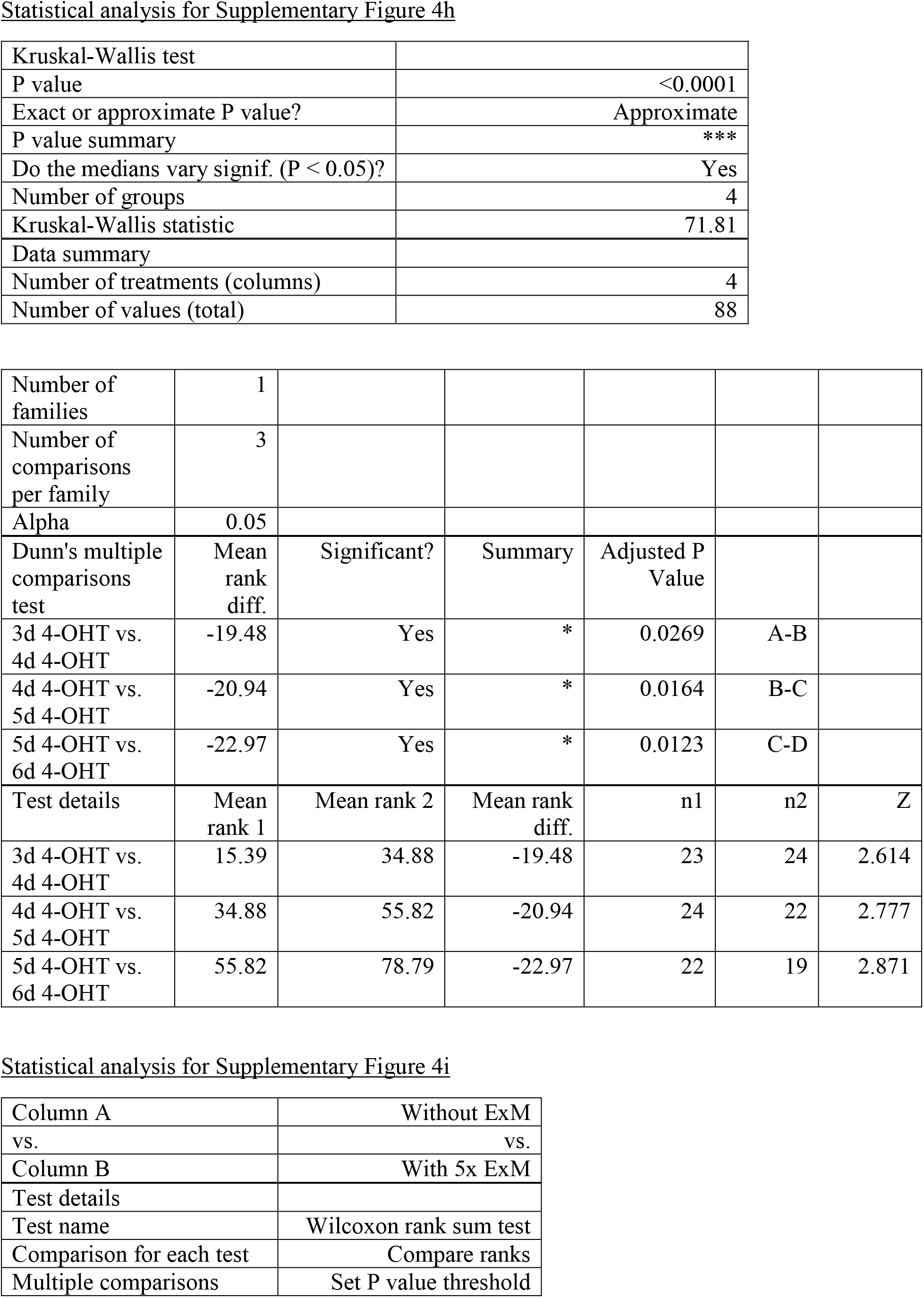
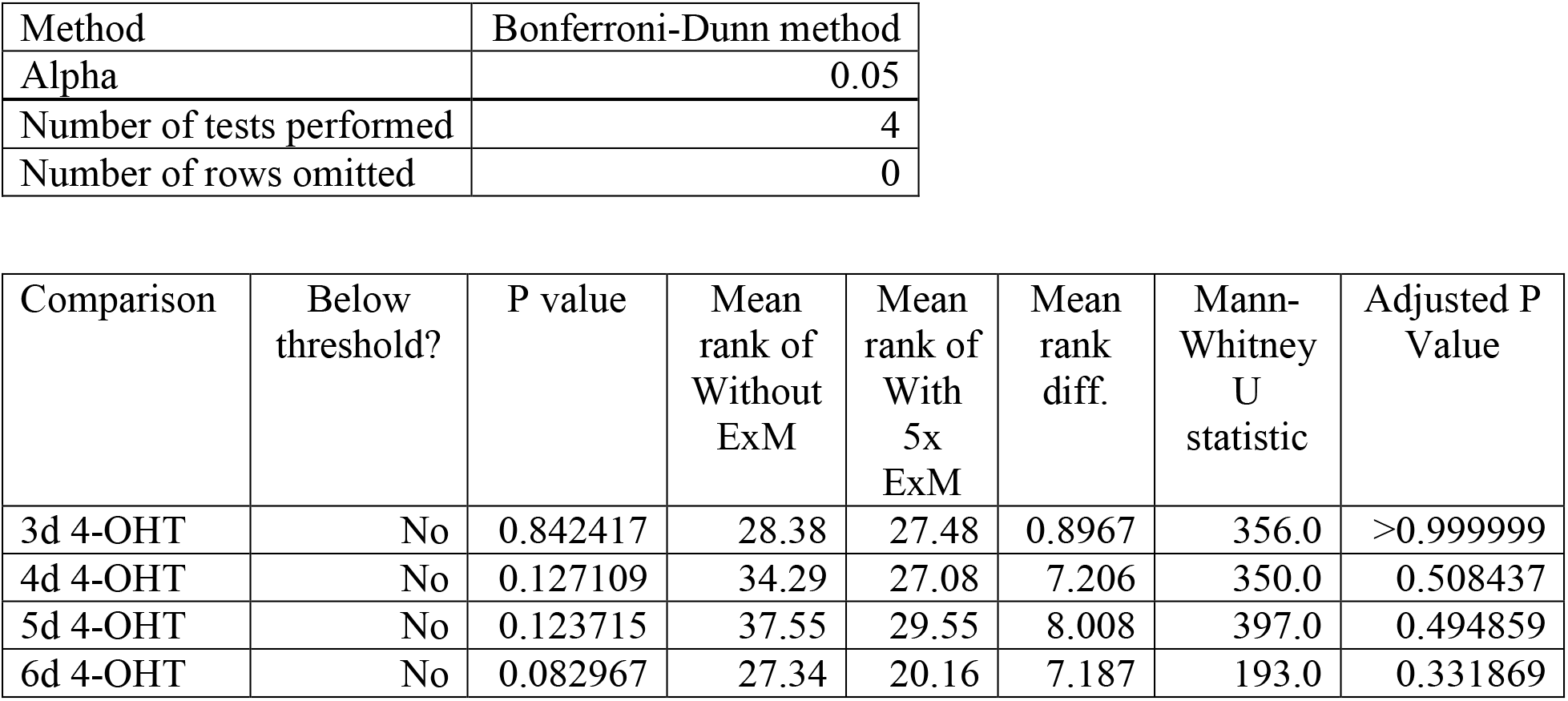
statistical analysis

Statistical analysis for Figure 2f
|  |  |
| --- | --- |
| Kruskal-Wallis test |  |
| P value | <0.0001 |
| Exact or approximate P value? | Approximate |
| P value summary | *** |
| Do the medians vary signif. (P < 0.05)? | Yes |
| Number of groups | 7 |
| Kruskal-Wallis statistic | 98.16 |
| Data summary |  |
| Number of treatments (columns) | 7 |
| Number of values (total) | 213 |

|  |  |  |  |  |  |  |
| --- | --- | --- | --- | --- | --- | --- |
| Number of families | 1 |  |  |  |  |  |
| Number of comparisons per family | 6 |  |  |  |  |  |
| Alpha | 0.05 |  |  |  |  |  |
| Dunn's multiple comparisons test | Mean rank diff. | Significant? | Summary | Adjusted P Value | A-? |  |
| No 4-OHT vs. 1d 4-OHT | -15.29 | No | ns | >0.9999 | B | 1d 4-OHT |
| No 4-OHT vs. 2d 4-OHT | -12.67 | No | ns | >0.9999 | C | 2d 4-OHT |
| No 4-OHT vs. 3d 4-OHT | -74.70 | Yes | ** | 0.0079 | D | 3d 4-OHT |
| No 4-OHT vs. 4d 4-OHT | -90.90 | Yes | *** | 0.0004 | E | 4d 4-OHT |
| No 4-OHT vs. 5d 4-OHT | -108.2 | Yes | *** | <0.0001 | F | 5d 4-OHT |
| No 4-OHT vs. 6d 4-OHT | -114.9 | Yes | *** | <0.0001 | G | 6d 4-OHT |

| Test details | Mean rank 1 | Mean rank 2 | Mean rank diff. | n1 | n2 | Z |
| --- | --- | --- | --- | --- | --- | --- |
| No 4-OHT vs. 1d 4-OHT | 36.33 | 51.62 | -15.29 | 9 | 21 | 0.6225 |
| No 4-OHT vs. 2d 4-OHT | 36.33 | 49.00 | -12.67 | 9 | 37 | 0.5530 |
| No 4-OHT vs. 3d 4-OHT | 36.33 | 111.0 | -74.70 | 9 | 32 | 3.212 |
| No 4-OHT vs. 4d 4-OHT | 36.33 | 127.2 | -90.90 | 9 | 38 | 3.979 |
| No 4-OHT vs. 5d 4-OHT | 36.33 | 144.6 | -108.2 | 9 | 47 | 4.826 |
| No 4-OHT vs. 6d 4-OHT | 36.33 | 151.2 | -114.9 | 9 | 29 | 4.885 |
| Statistical analysis for Figure 2m |  |
| Kruskal-Wallis test |  |
| P value | <0.0001 |
| Exact or approximate P value? | Approximate |
| P value summary | *** |
| Do the medians vary signif. (P < 0.05)? | Yes |
| Number of groups | 5 |
| Kruskal-Wallis statistic | 96.62 |
| Data summary |  |
| Number of treatments (columns) | 5 |
| Number of values (total) | 147 |

|  |  |  |  |  |  |  |
| --- | --- | --- | --- | --- | --- | --- |
| Number of families | 1 |  |  |  |  |  |
| Number of comparisons per family | 3 |  |  |  |  |  |
| Alpha | 0.05 |  |  |  |  |  |
| Dunn's multiple comparisons test | Mean rank diff. | Significant? | Summary | Adjusted P Value |  |  |
| 3d 4-OHT vs. 4d 4-OHT | -26.89 | Yes | * | 0.0255 | A-B |  |
| 4d 4-OHT vs. 5d 4-OHT | -28.30 | Yes | ** | 0.0069 | B-C |  |
| 5d 4-OHT vs. 6d 4-OHT | -44.80 | Yes | *** | <0.0001 | C-D |  |
| Test details | Mean rank 1 | Mean rank 2 | Mean rank diff. | n1 | n2 | Z |
| 3d 4-OHT vs.<br>4d 4-OHT | 28.88 | 55.76 | -26.89 | 32 | 38 | 2.632 |
| 4d 4-OHT vs.<br>5d 4-OHT | 55.76 | 84.06 | -28.30 | 38 | 47 | 3.047 |
| 5d 4-OHT vs.<br>6d 4-OHT | 84.06 | 128.9 | -44.80 | 47 | 29 | 4.456 |
| Statistical analysis for Figure 3f |  |
| Column B | KCl Stim |
| vs. | vs. |
| Column A | No Stim |
| Wilcoxon rank sum test |  |
| P value | <0.0001 |
| Exact or approximate P value? | Exact |
| P value summary | *** |
| Significantly different (P < 0.05)? | Yes |
| One- or two-tailed P value? | Two-tailed |
| Sum of ranks in column A,B | 528 , 507 |
| Mann-Whitney U statistic | 63 |
| Difference between medians |  |
| Median of column A | 0.002918, n=30 |
| Median of column B | 0.3804, n=15 |
| Difference: Actual | 0.3775 |
| Difference: Hodges-Lehmann | 0.3561 |
| Statistical analysis for Figure 4e |  |
| Column B | 10d 4-OHT |
| vs. |  |
| Column A | No 4-OHT |
| Wilcoxon rank sum test |  |
| P value | <0.0001 |
| Exact or approximate P value? | Approximate |
| P value summary | *** |
| Significantly different (P < 0.05)? | Yes |
| One- or two-tailed P value? | Two-tailed |
| Sum of ranks in column A,B | 242324, 869962 |
| Mann-Whitney U statistic | 63223 |
| Difference between medians |  |
| Median of column A | "0.003245, n=598" |
| Median of column B | "2.797, n=893" |
| Difference: Actual | 2.794 |
| Difference: Hodges-Lehmann | 2.778 |
| Statistical analysis for Supplementary Figure 4h |  |
| Kruskal-Wallis test |  |
| P value | <0.0001 |
| Exact or approximate P value? | Approximate |
| P value summary | *** |
| Do the medians vary signif. (P < 0.05)? | Yes |
| Number of groups | 4 |
| Kruskal-Wallis statistic | 71.81 |
| Data summary |  |
| Number of treatments (columns) | 4 |
| Number of values (total) | 88 |

|  |  |  |  |  |  |  |
| --- | --- | --- | --- | --- | --- | --- |
| Number of families | 1 |  |  |  |  |  |
| Number of comparisons per family | 3 |  |  |  |  |  |
| Alpha | 0.05 |  |  |  |  |  |
| Dunn's multiple comparisons test | Mean rank diff. | Significant? | Summary | Adjusted P Value |  |  |
| 3d 4-OHT vs. 4d 4-OHT | -19.48 | Yes | * | 0.0269 | A-B |  |
| 4d 4-OHT vs. 5d 4-OHT | -20.94 | Yes | * | 0.0164 | B-C |  |
| 5d 4-OHT vs. 6d 4-OHT | -22.97 | Yes | * | 0.0123 | C-D |  |
| Test details | Mean rank 1 | Mean rank 2 | Mean rank diff. | n1 | n2 | Z |
| 3d 4-OHT vs. 4d 4-OHT | 15.39 | 34.88 | -19.48 | 23 | 24 | 2.614 |
| 4d 4-OHT vs. 5d 4-OHT | 34.88 | 55.82 | -20.94 | 24 | 22 | 2.777 |
| 5d 4-OHT vs. 6d 4-OHT | 55.82 | 78.79 | -22.97 | 22 | 19 | 2.871 |
| Statistical analysis for Supplementary Figure 4i |  |
| Column A | Without ExM |
| vs. | vs. |
| Column B | With 5x ExM |
| Test details |  |
| Test name | Wilcoxon rank sum test |
| Comparison for each test | Compare ranks |
| Multiple comparisons | Set P value threshold |
| Method | Bonferroni-Dunn method |
| Alpha | 0.05 |
| Number of tests performed | 4 |
| Number of rows omitted | 0 |

| Comparison | Below threshold? | P value | Mean rank of Without ExM | Mean rank of With 5x ExM | Mean rank diff. | Mann-Whitney U statistic | Adjusted P Value |
| --- | --- | --- | --- | --- | --- | --- | --- |
| 3d 4-OHT | No | 0.842417 | 28.38 | 27.48 | 0.8967 | 356.0 | >0.999999 |
| 4d 4-OHT | No | 0.127109 | 34.29 | 27.08 | 7.206 | 350.0 | 0.508437 |
| 5d 4-OHT | No | 0.123715 | 37.55 | 29.55 | 8.008 | 397.0 | 0.494859 |
| 6d 4-OHT | No | 0.082967 | 27.34 | 20.16 | 7.187 | 193.0 | 0.331869 |

## Methods

### Animals and neuron cultures

All procedures involving animals at the Massachusetts Institute of Technology were conducted in accordance with the US National Institutes of Health Guide for the Care and Use of Laboratory Animals and approved by the Massachusetts Institute of Technology Committee on Animal Care and Biosafety Committee.

For **Figures 1, 2, 3** and **Supplementary Figures 1, 2, 3, 4**, hippocampal neurons were prepared from postnatal day 0 or 1 Swiss Webster mice (Taconic) (both male and female mice were used) as previously described^48^ with the following modifications: dissected hippocampal tissue was digested with 50 units of papain (Worthington Biochem) for 6-8 minutes, and the digestion was stopped with ovomucoid trypsin inhibitor (Worthington Biochem). Cells were plated at a density of 20,000-30,000 per glass coverslip coated with diluted Matrigel in a 24-well plate. Cells were seeded in 100 μL neuron culture media containing Minimum Essential Medium (MEM, no glutamine, no phenol red; Gibco), glucose (25 mM, Sigma), holo-Transferrin bovine (100 μg/mL, Sigma), HEPES (10 mM, Sigma), glutaGRO (2 mM, Corning), insulin (25 μg/mL, Sigma), B27 supplement (1X, Gibco), and heat inactivated fetal bovine serum (10% in volume, Corning), with final pH adjusted to 7.3-7.4 using NaOH. After cell adhesion, additional neuron culture media was added. AraC (2 μM, Sigma) was added at 2 days in vitro (DIV 2), when glial density was 50-70% of confluence. Neurons were grown at 37°C and 5% CO_2_ in a humidified atmosphere in a neuron incubator, with 2 ml total media volume in each well of the 24-well plate.

### Molecular Cloning

The DNAs encoding the protein motifs used in this study were mammalian-codon optimized and synthesized by Epoch Life Science and then cloned into mammalian expression backbones, pAAV-UBC (for constitutive expression), pAAV-UBC-FLEX (for Cre-dependent expression), or pAAV-cFos (for expression driven by the c-fos promoter) for DNA transfection in cultured neurons and AAV production by Janelia Viral Tools. See **Supplementary Table 1** for sequences of the motifs; see **Supplementary Table 2** for all tested constructs.

### DNA Transfection and AAV Transduction

For **Figure 1** and **Supplementary Figures 1** and **2**, cultured neurons were transfected at 7 days in vitro (DIV) with a commercial calcium phosphate transfection kit (Invitrogen) as previously described^49^. Briefly, for transfection in each coverslip/well in the 24-well plate, 5-50 ng of total XRI plasmid (5-25 ng of each plasmid when co-transfecting multiple plasmids), 200 ng pAAV-Syn-ERT2-iCre-ERT2 plasmid (only added in neurons for 4-OHT induction experiments), and pUC19 plasmid as a ‘dummy’ DNA plasmid to bring the total amount of DNA to 1500 ng (to avoid variation in DNA-calcium phosphate co-precipitate formation) were used. The cells were washed with acidic MEM buffer (containing 15 mM HEPES with final pH 6.7-6.8 adjusted with acetic acid (Millipore Sigma)) after 45-60 minutes of calcium phosphate precipitate incubation to remove residual precipitates.

For **Figures 2, 3** and **Supplementary Figures 3, 4**, cultured neurons were transduced at 7 days in vitro (DIV) with AAVs by adding the concentrated AAV stocks (serotype AAV9; Janelia Viral Tools) into neuron culture media at the following final concentrations in the 2 ml neuron culture media per well: for 4-OHT induction experiments, AAV9-UBC-XRI-HA at 5.56×10^9^ GC/ml, AAV9-UBC-FLEx-XRI-FLAG at 1.88×10^10^ GC/ml, and AAV9-Syn-ERT2-iCre-ERT2 at 8.60×10^9^ GC/ml; for c-fos recording experiments, AAV9-UBC-XRI-HA at 5.56×10^9^ GC/ml and AAV9-cFos-XRI-V5 at 1.39×10^9^ GC/ml.

### Chemical Treatments and Stimulations of Cultured Cells

For 4-OHT treatments in **Figures 1, 2** and **Supplementary Figures 2, 3, 4**, the original culture media of neuron cultures was transferred into a new 24-well plate, where the media from different neuron cultures were stored in different wells, and kept in the neuron incubator until the end of 4-OHT treatment. 2 ml of fresh neuron culture media containing 1 μM 4-hydroxytamoxifen (4-OHT; Sigma H6278) was added into each well of neuron culture. The neuron cultures were then placed back to the neuron incubator and incubated for 15 minutes, followed by a brief wash in MEM media. Finally, the MEM media was removed and the original neuron culture media was transferred back to the corresponding wells of neuron culture. The neuron cultures were then placed back into the neuron incubator.

For potassium chloride (KCl) treatments in **Figure 3**, the KCl depolarization solution was prepared, which contained 170 mM KCl, 2 mM CaCl_2_, 1 mM MgCl_2_, and 10 mM HEPES. Then, KCl depolarization media was prepared by mixing KCl depolarization solution and fresh neuron culture media at the volume ratio of 1: 2.32, so that the final concentration of K^+^ after mixing was 55 mM (taking account the K^+^ from the fresh neuron culture media). The original culture media of neuron cultures was transferred into a new 24-well plate, where the media from different neuron cultures were stored in different wells, and kept in the neuron incubator until the end of the KCl-induced depolarization treatment. 2 ml of KCl depolarization media was added to each well of neuron culture. Neuron cultures were then placed back into the neuron incubator and incubated for 3 hours. Finally, the KCl depolarization media was removed and the original neuron culture media was transferred back into the corresponding wells of the neuron cultures. The neuron cultures were then placed back to the neuron incubator.

### Animals and mouse surgery

All procedures involving animals at Boston University were conducted in accordance with the US National Institutes of Health Guide for the Care and Use of Laboratory Animals and approved by the Boston University Institutional Animal Care and Use and Biosafety Committees.

For experiments in **Figure 4**, all surgeries were performed under stereotaxic guidance, and coordinates were given relative to bregma (in mm). Dorsal ventral injections were calculated and zeroed out relative to the skull. Wild type C57BL/6 mice (3 months of age; male; Charles River Labs) were placed into a stereotaxic frame (Kopf Instruments) and anesthetized with 3% isoflurane during induction (lowered to 1-2% to maintain anesthesia throughout the surgery). Ophthalmic ointment was applied to both eyes. Hair was removed with a hair removal cream and the surgical site was cleaned with ethanol and betadine. Following this, an incision was made to expose the skull. Bilateral craniotomies involved drilling windows through the skull above the injection site using a 0.5 mm diameter drill bit. Coordinates were −2.0 anteroposterior (AP), ±1.5 mediolateral (ML), and −1.5 dorsoventral (DV) for dorsal CA1.

The AAV mixture for injection was prepared by mixing the AAV stocks (serotype AAV9; Janelia Viral Tools) at the following final concentrations: AAV9-UBC-XRI-HA at 1.48×10^13^ GC/ml, AAV9-UBC-FLEx-XRI-FLAG at 3.77×10^13^ GC/ml, and AAV9-Syn-ERT2-iCre-ERT2 at 1.72×10^13^ GC/ml. Mice were injected with 1 μl of the AAV mixture at the target site using a mineral oil-filled 33-gauge beveled needle attached to a 10 μl Hamilton microsyringe (701LT; Hamilton) in a microsyringe pump (UMP3; WPI). The needle remained at the target site for five minutes post-injection before removal. Mice received buprenorphine intraperitoneally following surgery and were placed on a heating pad during surgery and recovery.

### 4-Hydroxytamoxifen injection

For experiments in **Figure 4**, 4-OHT (Sigma) was dissolved in 100% ethanol (Sigma) at 100 mg/ml by vortexing for 5 minutes. Next, the solution was mixed with corn oil (Sigma) to obtain a final concentration of 10 mg/ml 4-OHT by vortexing for 5 minutes and then sonicating for 30-60 minutes until the solution was clear. The 10 mg/ml 4-OHT solution was then loaded into syringes and administered to mice via intraperitoneal (i.p.) injection at 40 mg/kg.

### Histology

For experiments in **Figure 4**, mice were transcardially perfused with 1X PBS followed by 4% paraformaldehyde in 1X PBS. The brain was gently extracted from the skull and post-fixed in 4% paraformaldehyde in 1X PBS overnight at 4°C. The brain was then incubated in 100 mM glycine in 1X PBS for 1 hour at RT, and then the brain was transferred into 1X PBS and stored at 4°C until slicing. The brain was sliced to 50-μm thickness coronally using a vibratome (Leica), and then stored in 1X PBS at 4°C until immunofluorescence staining.

### Immunofluorescence

#### Immunofluorescence of cultured cells

**Figures 1, 2, 3** and **Supplementary Figures 1, 2, 3**, cells were fixed in TissuePrep buffered 10% formalin for 10 minutes at room temperature (RT) followed by three washes in 1X PBS, 5 minutes each at RT. Cells were then incubated in MAXBlock Blocking medium (Active Motif) supplemented with final concentrations of 0.1% Triton X-100 and 100 mM glycine for 20 minutes at RT, followed by three washes in MAXwash Washing Medium (Active Motif), 5 minutes each at RT. Next, cells were incubated with primary antibodies in MAXStain Staining medium (Active Motif) at 1:500 overnight at 4°C, followed by three washes in MAXwash Washing medium, 5 minutes each at RT. Cells were then incubated with fluorescently-labeled secondary antibodies and NeuroTrace Blue Fluorescent Nissl Stain (Invitrogen) in MAXStain Staining medium, all at 1:500, overnight at 4°C, followed by three washes in MAXwash Washing medium, 5 minutes each at RT. The cells were then stored in 1X PBS at 4°C until imaging.

#### Immunofluorescence of brain slices

For **Figure 4**, brain slices were blocked overnight at 4°C in MAXBlock Blocking medium, followed by four washes for 30 minutes each at RT in MAXWash Washing medium. Next, slices were incubated with primary antibodies in MAXStain Staining medium at 1:250 overnight at 4°C, and then washed in MAXWash Washing medium four times for 30 minutes each at RT. Next, slices were incubated with fluorescently-labeled secondary antibodies at 1:500 and NeuroTrace Blue Fluorescent Nissl Stain (Invitrogen) at 1:250 in MAXStain Staining medium overnight at 4°C, and then washed in MAXWash Washing medium four times for 15 minutes each at RT. The slices were then stored in 1X PBS at 4°C until imaging.

#### Expansion microscopy of cultured cells

For **Supplementary Figure 4**, cell cultures on round coverslips were fixed in 4% paraformaldehyde (Electron Microscopy Sciences) and 0.1 % glutaraldehyde (Electron Microscopy Sciences) in 1X PBS for 10 min at RT. Cells were then incubated in 0.1 % sodium borohydride (Sigma) in 1X PBS for 7 min and then 100 mM glycine (Sigma) in 1X PBS for 10 min, both at RT.

Acryloyl-X (6-((acryloyl)amino)hexanoic acid, succinimidyl ester (AcX) (Invitrogen) was resuspended in anhydrous DMSO (Invitrogen) at a concentration of 10 mg/ml, and stored in a desiccated environment at −20°C. For anchoring, cells were incubated in 200 μL of AcX at a concentration of 0.1 mg/ml in a 2-(N-morpholino)ethanesulfonic acid (MES)-based saline (100 mM MES, 150 mM NaCl) overnight at 4°C. Then, cells were washed with 1X PBS three times at RT for 5 minutes each.

Gelation solution which contains 1.1 M sodium acrylate (Sigma), 2 M acrylamide (Sigma), 90 ppm N,N’-methylenebisacrylamide (Sigma), 1.5 ppt ammonium persulfate (APS) (Sigma), and 1.5 ppt tetramethylethylenediamine (TEMED) (Sigma) in 1X PBS was prepared fresh. Cells were first incubated on ice for 10 min with shaking to prevent premature gelation and enable diffusion of solution into samples. A gelation chamber was prepared by placing two No. 1.5 coverslips on a glass slide spaced by about 8 mm to function as insulators on either end of the neuronal coverslip to avoid compression and each coverslip containing a neuronal cell culture sample was placed on a gelation chamber with the cells facing down. The gelation chamber was filled with gelation solution and a coverslip placed over the sample and across the two insulators to ensure the sample was covered with gelling solution and no air bubbles were formed on the sample. Samples incubated at 37°C for 1 hours in a humidified atmosphere to complete gelation. Following gelation, the top coverslip was removed from the samples, and only the sample gel was transferred into a 1.5 mL tube containing 1 mL of denaturation buffer, consisting of 5% (w/v) sodium dodecyl sulfate (SDS), 200 mM NaCl, and 50 mM Tris at pH 8. Gels were incubated in denaturation buffer overnight at RT and 3 hour at 80°C, followed by washing in water overnight at RT to remove residual SDS. Gels were then stored in 1X PBS at 4°C before immunostaining.

For immunostaining and imaging, gels were first incubated in bovine serum albumin (BSA) blocking solution that contains 1% BSA, 0.5% Triton-X in 1X PBS for 1 hour at RT then with primary antibodies overnight at 4°C. Gels were washed three times in BSA blocking solution for 30 minutes each at RT and incubated with fluorescently-labeled secondary antibodies overnight at 4°C. Gels were then washed three times in BSA blocking solution for 30 minutes each at RT and expanded in water overnight at 4°C before imaging.

#### Antibodies and Nissl Stain

The following antibodies and Nissl stain were used in this paper: primary antibodies, anti-HA (Santa Cruz, cat# sc-7392), anti-FLAG (Invitrogen, cat# 740001), anti-V5 (Abcam, cat# ab9113), anti-NeuN (Synaptic Systems, cat# 266004); fluorescent secondary antibodies from Invitrogen, cat# A-21241, cat# A-21133, cat# A-32933, cat# A-32733, cat# A-11035, and cat# A-11073; fluorescent secondary antibodies from Biotium, cat# 20308; Nissl stain, NeuroTrace Blue Fluorescent Nissl Stain (Invitrogen, cat# N21479).

### Fluorescence Microscopy of Immunostained Samples

Fluorescence microscopy was performed on a spinning disk confocal microscope (a Yokogawa CSU-W1 Confocal Scanner Unit on a Nikon Eclipse Ti microscope) equipped with a 40X 1.15 NA water immersion objective (Nikon MRD77410), a Zyla PLUS 4.2 Megapixel camera controlled by NIS-Elements AR software, and laser/filter sets for 405 nm, 488 nm, 561 nm, and 640 nm optical channels. For each field of view, multi-channel volumetric imaging was performed at 0.4 μm per Z step. Imaging parameters were kept the same for all samples within a set of experiments (e.g., a set of 4-OHT induction experiments in which samples were treated with 4-OHT at different time points).

### Image Analysis

Image analysis was performed in ImageJ (ImageJ National Institutes of Health) and MATLAB (MathWorks).

#### Intensity profile measurements

First, the somata of neurons in the images were identified by the Nissl staining (in samples without ExM) or anti-NeuN staining (in samples with ExM) channel, and XRI(s) in the soma of each neuron were identified by the anti-HA channel. If multiple XRIs were present in a soma, the XRI with the longest length as well as any XRI with length above half of that longest length was selected for downstream analysis. For each XRI, a curved centerline was drawn along the longitudinal direction of XRI in the anti-HA channel. The centerline width was set to half of the width of the XRI. The intensity profiles along this centerline were measured in the anti-HA channel (and called the HA line profile) and in other XRI epitope staining channels, such as in the anti-FLAG channel (called the FLAG line profile) or anti-V5 channel (called the V5 line profile).

#### Readout information from intensity profiles

See **Supplementary Figure 2** for the process flow of extracting information from the intensity profiles of XRIs. **Step 1**: For each XRI, a curved centerline was drawn along the longitudinal axis of the XRI in the anti-HA channel. The centerline width was set to half of the width of the XRI. **Step 2**: The intensity profiles along this centerline were measured in the anti-HA channel (resulting in an HA line profile) and in the other XRI epitope staining channel, such as in the anti-FLAG channel (resulting in a FLAG line profile) or in the anti-V5 channel (resulting in a V5 line profile). **Step 3**: Next, each of the line profiles was split into two half line profiles using the geometric center point of the XRI (the 50% length point along the centerline, measuring from the end of the XRI) as the ‘split point’. Each of the half HA line profiles was then converted into a line integrals of HA, by integrating the line profile with respect to the distance along the half centerline starting from the geometric center point, and then these line integrals of HA were normalized to the maximum integral value so that each line integral of HA started at the value 0 at the geometric center point of the XRI, and gradually increased to the value 1 at the end of the XRI. For the corresponding half FLAG (or V5) line profiles, line integrals were also calculated but not normalized. At this point, we have the line integrals of HA and FLAG (or V5), which correspond to the cumulative HA and FLAG (or V5) intensities along each half of the XRI. The FLAG (or V5) intensity change per unit change in the cumulative HA intensity, defined as the FLAG (or V5) signal, was calculated by taking the derivative of the line integral of FLAG (or V5) with respect to the line integral of HA. At this stage, we obtained the line integral of HA and the FLAG (or V5) signal from each of the halves of the XRI, and the final extracted FLAG (or V5) signal from this XRI was defined as the point-by-point average of the two FLAG (or V5) signals from the two halves of the XRI. **Step 4**: We found the two obtained FLAG (or V5) signals from the same XRI have small but noticeable differences. We reasoned that such small but noticeable discrepancies between the two halves of the same XRI was due to the asymmetry of the XRI, and the choice of the exact geometric center as the split point may not be optimal. To minimize the discrepancy between the two FLAG (or V5) signals from the two halves of the same XRI, we searched for an optimal split point near the geometric center of the XRI (searching range was the geometric center +/- 10% of the total XRI length), so that using this optimal split point, instead of the geometric center, as the split point results in the least difference (in sum of squared differences) between the two FLAG (or V5) signals from the two halves of the splitted XRI. **Step 5**: Same as Step 3, except that the optimal split point, instead of the geometric center, was used to split the intensity profiles into two halves. We found the resulting final FLAG (or V5) signal (after averaging those from the two halves) when using the geometric center as the split point was similar to that when using the optimal split point as the split point. Nevertheless, we used the optimal split point as the split point to analyze XRIs throughout this paper.

#### Calculation of the fraction of HA line integral when FLAG signal begins to rise

The FLAG signal minus the FLAG signal at the center of XRI (i.e., the optimal split point as defined above) was plotted against the fraction of HA line integral. The initial rising phase of the FLAG signal (defined as the portion of the FLAG signal between 10% to 50% of the peak FLAG signal) was fitted as a linear function, which was then extrapolated onto the axis of the fraction of HA line integral. The intersection point at the axis of the fraction of the HA line integral was defined as the fraction of HA line integral when the FLAG signal began to rise.

### Statistical analysis

All statistical analysis was performed using the built-in statistical analysis tools in Prism (GraphPad) or MATLAB. The statistical details of each statistical analysis can be found in the figure legends and **Supplementary Table 3**.

